# Microbiome composition shapes temperature tolerance in a Hawaiian picture-winged *Drosophila*

**DOI:** 10.1101/2025.06.03.657679

**Authors:** Donald K. Price, Kristian West, Michelle Cevallos-Zea, Sara Helms Cahan, Joaquin C. B. Nunez, Emily K. Longman, Joanne Y. Yew, Matthew J. Mederios

## Abstract

Hawaiian picture-winged *Drosophila* are undergoing rapid biodiversity loss, with twelve species listed as endangered and others in decline. Gut microbiota are increasingly recognized as contributors to host adaptation that are capable of influencing stress tolerance, reproduction, and other fitness-related traits. We investigated the role of microbial communities in local adaptation using two populations of *Drosophila basisetae* from Hawaiian rainforests at 900 m and 1200 m elevation. Microbiome profiling of wild flies by high throughput amplicon sequencing revealed distinct bacterial and fungal communities between sites. Whole-genome resequencing of the two *Drosophila* populations identified weak but significant population genetic structure, with evidence of admixture and gene flow. Despite this connectivity, 16 outlier SNPs across 18 genes showed patterns consistent with divergent selection, suggesting localized adaptation. To test microbiome effects on host physiology experimentally, we conducted a fully factorial research design with microbiome inoculations in laboratory-reared flies acclimated to 18 °C (control) or 24 °C (stressful). Flies treated with low-elevation microbiota had higher survival across temperatures, whereas those treated with high-elevation microbiota produced more eggs, indicating microbiome-mediated differences in reproductive investment. Activity levels at 18 °C were higher when flies received microbiota from their native population. Measures of critical thermal maximum (CTmax) and male accessory gland size showed complex interactions among microbiome source, temperature, and fly population. These results indicate that microbes may modulate host thermal tolerance and reproduction in environmentally-dependent and population-specific ways. Our findings suggest that microbiome-host-environment interactions may contribute to both phenotypic plasticity and evolutionary adaptation to enhance resilience to environmental stress, with important implications for conservation in rapidly changing Hawaiian ecosystems.

**Summary statement:** This study demonstrates that microbiome composition influences thermal tolerance through alterations in survival and reproduction in the Hawaiian picture wing, *Drosophila basisetae*, potentially contributing to adaptive phenotypic changes between populations.

## INTRODUCTION

The gut microbiome can play an important role in shaping host physiology and fitness across a wide range of animals, including insects. In *Drosophila*, bacterial taxa break down simple and complex carbohydrates while modulating host lipid and carbohydrate metabolism contributing to digestion, metabolic regulation, immune function, and reproductive success (Adair et al., 2018; Broderick & Lemaitre, 2012; Gould et al., 2018). Fungal taxa contribute to the degradation of complex polymers and may help stabilize gut conditions during stress (Medeiros et al., 2024; Nash et al., 2017; Suen et al., 2010; Walters et al., 2020). Together, these microbial communities mediate host responses to environmental variability through immediate physiological adjustments and longer-term developmental phenotypic plasticity (Henry et al., 2013; McMullen et al., 2020; Moghadam et al., 2018; Shin et al., 2011; Téfit et al., 2023; Wong et al., 2013). Environmental conditions, such as temperature, humidity, and diet, strongly influence microbiome composition, creating context-dependent relationships between microbial taxa and host traits (Agler et al., 2016; Douglas, 2015). Importantly, the effects of the microbiome are not fixed; they can vary across host genotypes, developmental stages, and environmental contexts, resulting in complex outcomes for stress tolerance, behavior, and reproduction (Amend et al., 2022; McMullen et al., 2020; Walters et al., 2020).

The microbiome’s ability to influence host phenotypes under variable conditions may be an important factor in local adaptation of host species. Recent studies have proposed that microbiome plasticity can enable organisms to cope with novel environments, potentially complementing or interacting with host genetic adaptation (Amend et al., 2022; Baldassarre et al., 2022). For example, artificial selection on thermal stress tolerance and lifespan in *Drosophila melanogaster* led to persistent divergence in gut microbiome composition among selected lines, even when reared under common conditions, highlighting the potential for host genomic background to shape microbial communities (Kristensen et al., 2024). Thus, variation in microbiome composition across habitats and populations may reflect adaptive processes shaped by ecological and evolutionary pressures. These insights are particularly relevant in spatially structured and environmentally heterogeneous systems, where both the host and its associated microbiota may undergo local specialization.

The Hawaiian Islands provide an ideal setting to investigate such dynamics. Hawaiian *Drosophila*, a classic example of adaptive radiation, occupy ecologically diverse forest habitats along sharp environmental gradients (Eldon et al., 2013; Magnacca & Price, 2015). These populations experience distinct environment conditions over short spatial scales, providing a natural system in which to explore how microbiomes and host genotypes interact with environmental variables. Previous work on Hawaiian arthropods has shown that gut microbial communities can vary by habitat, sex, and diet, and may influence reproductive success and other fitness-related traits (Armstrong et al., 2022; Poff et al., 2017; Téfit et al., 2023). Research on Hawaiian *Drosophila* shows that bacterial and fungal components of the gut microbiome can have distinct, sex-specific roles in shaping reproductive success (Medeiros et al., 2024). Such findings suggest that both microbiome composition and its functional consequences may be shaped by local ecological conditions and evolutionary histories.

This study focuses on the microbiome of *Drosophila basisetae*, an endemic Hawaiian picture-wing *Drosophila* whose relatives face severe conservation challenges (Corpuz et al., 2023; Magnacca & Price, 2015) with 12 species listed as endangered species (U.S. Fish & Wildlife Service, 2006), and is structured around two general multifaceted hypotheses. First, we propose that environmental variation shapes host-associated microbial composition, with flies from distinct microclimates hosting different communities (Agler et al., 2016; Zhao et al., 2023) and that the differences in the microbiomes significantly influence host fitness traits, including thermal tolerance, reproductive investment, and stress resilience, in a manner that contributes to local adaptation (Amend et al., 2022; Gould et al., 2018; McMullen et al., 2020; Medeiros et al., 2024; Téfit et al., 2023; Walters et al., 2020). Second, more broadly we propose that complex interactions among microbiome origin, host genetics, and acclimation environment will generate variable phenotypic outcomes, driving local genetic divergence and contributing to adaptive evolution (Amend et al., 2022; Téfit et al., 2023). By integrating microbiome inoculation, thermal stress physiology, and host genomic analyses across contrasting environments, this work aims to elucidate how microbiome–host– environment interactions shape phenotypic plasticity and adaptation in natural populations, thereby advancing our understanding of ecological adaptation and evolutionary processes (Kang et al., 2017, 2016). In doing so, our ongoing research on the Hawaiian *Drosophila* contributes to a growing body of research exploring the role of the microbiome in ecological adaptation and evolutionary processes.

## MATERIALS AND METHODS

### Fly populations: Wild fly collections

*D. basisetae* flies were collected from the wild at two East Hawaiʻi Island sites located approximately 13 km apart and differing by 237 m in elevation (**Table 1**). A total of 45 adult *D. basisetae* from Ola’a and 52 adult *D. basisetae* from Tom’s Trail were captured during two collection trips (August 2021 and February 2022) and brought back to the laboratory at the University of Nevada, Las Vegas (UNLV) under NPS permits HAVO-2021-SCI-0013 & HAVO-2024-SCI-0028 and State of Hawaii permits I2978 & I5297 (**Fig. 1**). Adult flies were collected using fermented banana and mushroom smeared on sponge baits: baits were checked every 15 minutes and flies were captured using large polypropylene tubes (35 mm x 100mm) and then aspirated into adult food vials in the field. All the collected flies were used to establish the laboratory populations were kept in adult food vials (see recipe below) that were placed in insulated containers with ice packs keeping temperature at 15-18 °C, until transported to UNLV 3-5 days later. The higher elevation site was in the Ola’a of the Hawaiʻi Volcanoes National Park at 1212 m elevation adjacent to Wright Road. The lower elevation site was in the Waiakea State Forest Reserve at an elevation of 975 m along Tom’s Trail approximately 580 m north of Stainback Hwy (**Table 1**). The temperature, rainfall and humidity data obtained from the Hawaiʻi Climate Data Portal for all years in the publicly available data and for the year 2022 (Longman et al., 2024). Local measurements were also taken during the times when the *D. basisetae* were collected for this study: Ola’a: 20 August 2021: 8-21 °C, 58-99 % relative humidity (RH), and 8 February 2022: 8-16 °C, 91-94 % RH; Tom’s Trail: 20 August 2021: 13-22 °C, 78-99 % RH, and 8 February 2022: 13-20 °C, 82-98 % RH.

**Fig 1.**
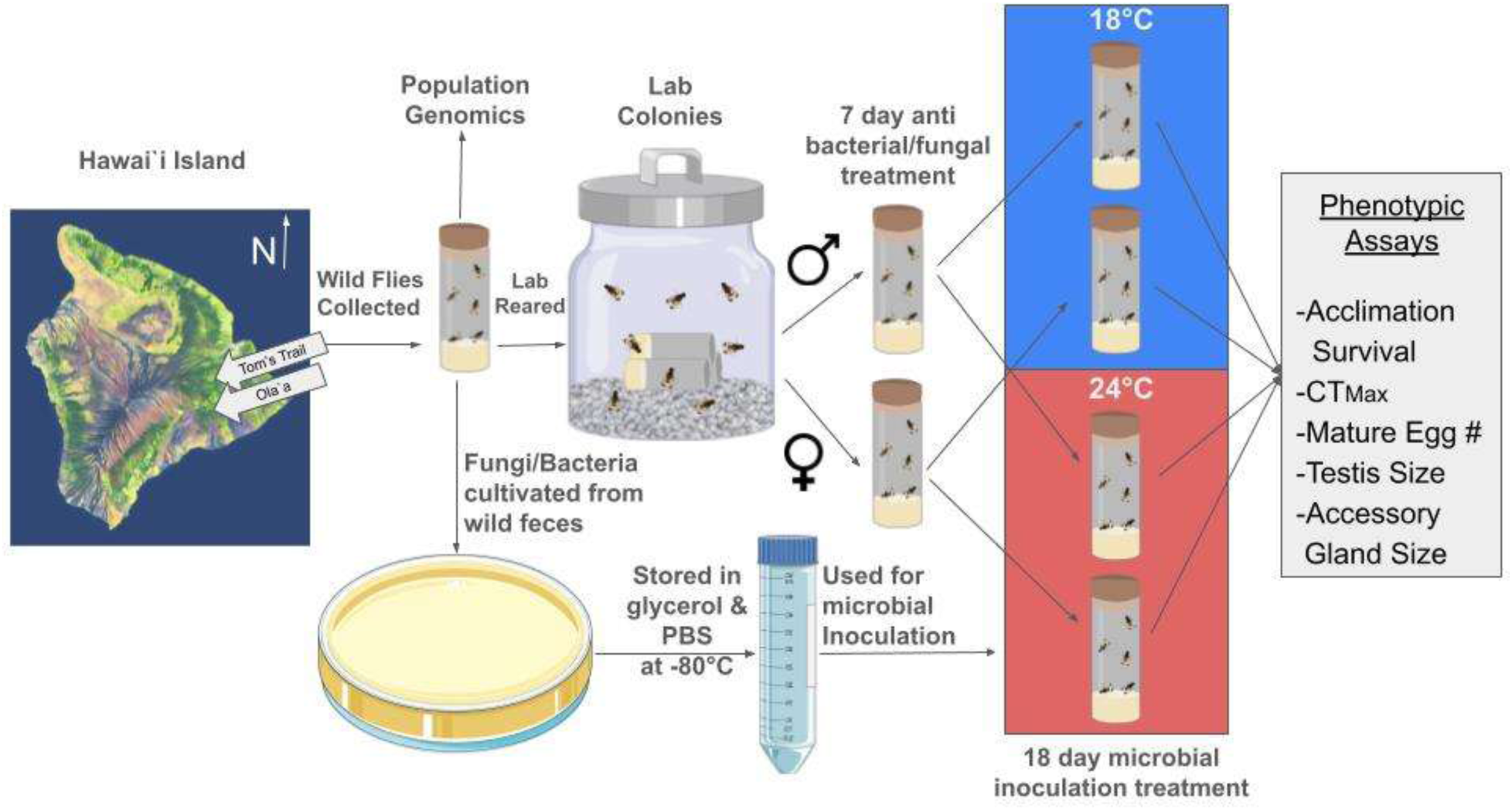
Schematic of the collection and raising of *Drosophila basisetae* populations including the experimental design, phenotypic assays, and population genomic analyses. The 18 °C and 24 °C temperatures are standard temperature and stressful temperatures for Hawaiian picture wing *Drosophila*, respectively (Eldon et al., 2019; Uy et al., 2015).

**Table 1.**
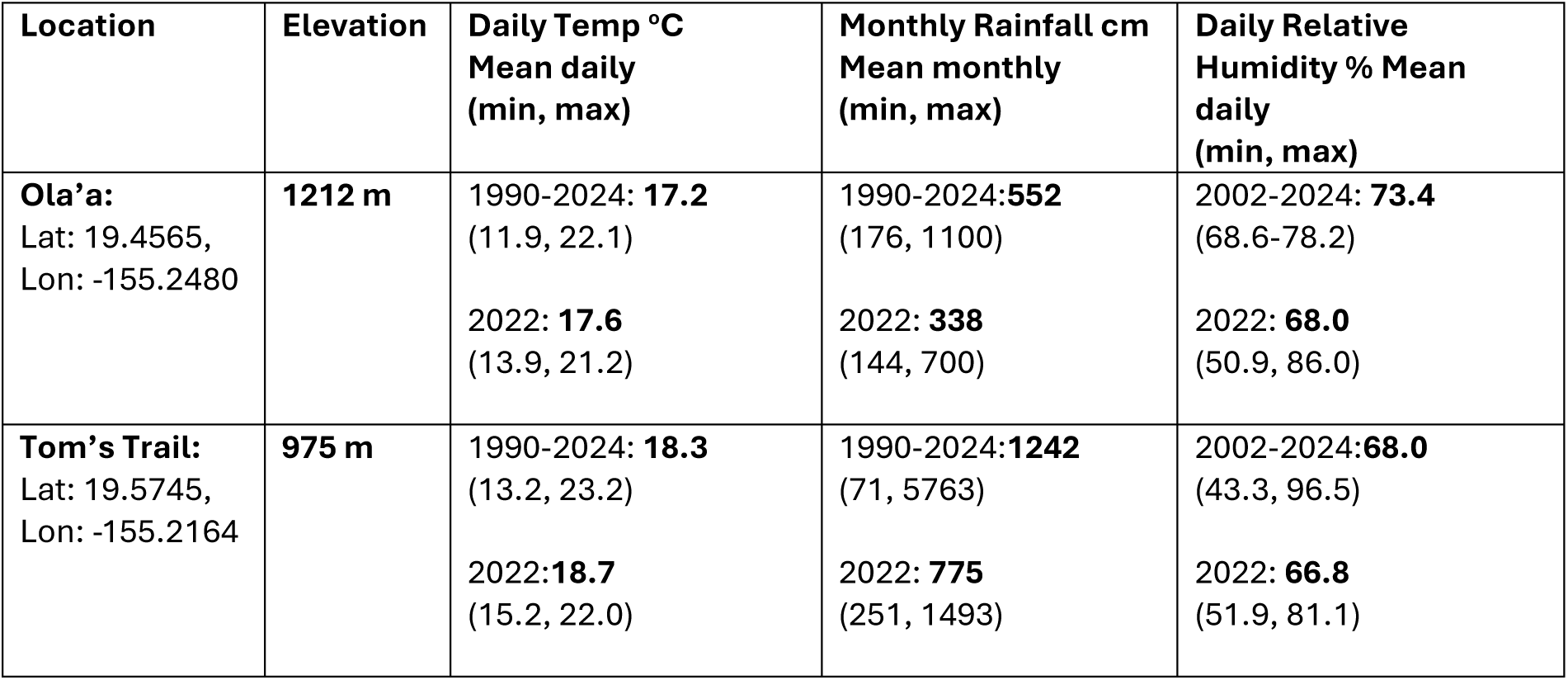
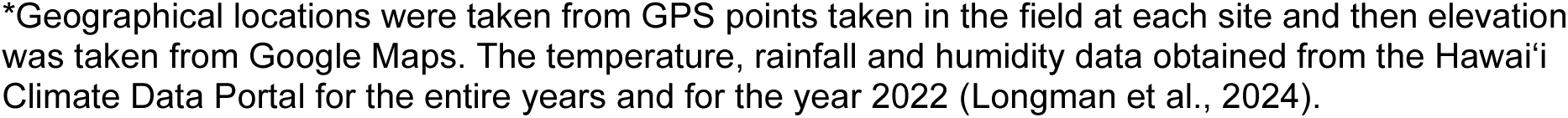
Climatic description of the two study locations on Hawaii Island: Ola’a in Volcanoes National Park, and Tom’s Trail in Waiakea State Forest Reserve*.

| Location | Elevation | Daily Temp °C<br>Mean daily<br>(min, max) | Monthly Rainfall cm<br>Mean monthly<br>(min, max) | Daily Relative<br>Humidity % Mean<br>daily<br>(min, max) |
| --- | --- | --- | --- | --- |
| **Ola'a:**<br>Lat: 19.4565,<br>Lon: -155.2480 | **1212 m** | 1990-2024: **17.2**<br>(11.9, 22.1)<br>2022: **17.6**<br>(13.9, 21.2) | 1990-2024:**552**<br>(176, 1100)<br>2022: **338**<br>(144, 700) | 2002-2024: **73.4**<br>(68.6-78.2)<br>2022: **68.0**<br>(50.9, 86.0) |
| **Tom's Trail:**<br>Lat: 19.5745,<br>Lon: -155.2164 | **975 m** | 1990-2024: **18.3**<br>(13.2, 23.2)<br>2022:**18.7**<br>(15.2, 22.0) | 1990-2024:**1242**<br>(71, 5763)<br>2022: **775**<br>(251, 1493) | 2002-2024:**68.0**<br>(43.3, 96.5)<br>2022: **66.8**<br>(51.9, 81.1) |

\*Geographical locations were taken from GPS points taken in the field at each site and then elevation was taken from Google Maps. The temperature, rainfall and humidity data obtained from the Hawai'i Climate Data Portal for the entire years and for the year 2022 (Longman et al., 2024).

### Population Genomic Analyses of wild collected Flies

We sequenced 22 wild-caught *D. basisetae* samples (7 from Tom’s Trail and 15 from Ola’a) from the wild flies that were used to establish the laboratory populations (**Fig. 1**). For these samples, DNA was extracted using the Zymo Quick-DNA Tissue/Insect Kit (D6017, Zymo Research Corporation, CA). Genomic libraries were prepared using a plexWell LP384 preparation kit (seqWell Inc., Beverly, MA) and sequenced in a NovaSeq machine in a 2x150 pair-end configuration by Novogene Corp to an approximate genome-wide mean coverage of 6X. Samples with mean coverage less than 6X were resequenced on a Singular Genomics G4 sequencer at the Vermont Integrated Genomics Core (VIGR) at the University of Vermont. We performed read quality control with FastQC (http://www.bioinformatics.babraham.ac.uk/projects/fastqc/) and MultiQC (Ewels et al., 2016). Reads were trimmed using Fast (Chen et al., 2018) and mapped to the *D. basisetae* reference genome (GenBank accession: GCA_035041595.1; (de Vries et al., 2017). Prior to mapping, we masked repetitive and low-complexity regions of the genome using RepeatMasker (using the model trained in *D. melanogaster*). Mapping was done using BWA-MEM2 (Jung & Han, 2022). Post-processing, including duplicate removal, was done using Picard [v3.1.1] and SAMtools [v1.9] (Li et al., 2009). Four individuals that had unusually low coverage (< 2.5X) were removed prior to single nucleotide polymorphism (SNP) calling (i.e., 18 individuals were used for subsequent steps).

The 18 individuals with sufficient coverage were genotyped using a two-pronged approach. First, we used a haplotype-based GATK v4.6 (McKenna et al., 2010) pipeline with stringent filtering parameters. Accordingly, BAM files generated with BWA-MEM2 were individually haplotyped assuming a ploidy of 2, a heterozygosity prior of 0.005, and using the GVCF as the mode for emitting reference confidence scores (i.e., the -ERC flag). We used these haplotype files to generate a joint GATK genomic database object and subsequently a VCF with genotype calls. The resulting VCF was further filtered to retain only bi-allelic sites with an average missing rate of 5% or less, a global allele frequency of 5% or more, and a minimum coverage of 4X across all individuals. Given that a genome-wide coverage of 4X can be considered “shallow” sequencing, we also conducted a parallel analysis using ANGSD v0.933 (Korneliussen et al., 2014) to estimate genotype likelihoods. For this analyses, we ran ANGSD using the GATK model (i.e., -GL 2) using the following parameters: -remove_bads 1 -baq 1 -skipTriallelic 1 -uniqueOnly 1 -only_proper_pairs 1 -C 50 -SNP_pval 1x10-6 - minMapQ 30 -minQ 20 -minMaf 0.1. During SNP calling, we filtered to only include SNPs that were present in 94.4% of individuals and had a minimum individual sequencing depth of 4X.

***Population Structure Analyses:*** We conducted parallel population structure analyses using data generated by both approaches. For the GATK haplotype-based approach, we estimated the fixation index (*F*_ST_) and kinship analyses in SNPRelate v3.19 (Zheng et al., 2012). We report between-family kinship using the KING-robust method (Manichaikul et al., 2010). For *F*_ST_, we used the Weir and Cockerham method (Weir & Cockerham, 1984). We also estimated principal component analysis (PCA), using FactoMineR v2.11 (Lê et al., 2008), and discriminant analyses of principal component (DAPC) using adegenet v2.1.10 (Jombart, 2008). For the DAPC analysis, we identified the optimal number of PCs to retain by using the *optim.a.score()* function. For the ANGSD-based analyses, *F*_ST_ was calculated using two-dimensional site frequency spectra (2dSFSs). Given unequal sample sizes between the two populations, we performed a bootstrap approach with 100 iterations. We also used PCAngsd v1.35 (Meisner & Albrechtsen, 2018) to estimate the covariance matrix and conduct a PCA using the BEAGLE file output from ANGSD.

***Local Adaptation Analyses:*** To test for signatures of local adaptation between our populations, we employed a generalized linear model (GLM) approach using a binomial error model. Specifically, we modeled the alternative allele proportions at each SNP that passed our filtering criteria as a function of two covariates: (1) the sample’s projection in PC1 space, which accounts for population structure, and (2) the population label (i.e., Ola‘a vs. Tom’s Trail) as a categorical factor. We fitted two models with a binomial error structure: the first included only the PC1 projections (Eq. 1), while the second incorporated both the PC1 projections and population labels (Eq. 2). To assess whether including the population label improved model fit, we performed a likelihood ratio test (LRT), comparing the model with both PC1 projection and habitat label—used as a proxy for potentially adaptive alleles—against the model with only PC1 projection, which accounts solely for population structure.

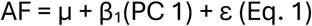

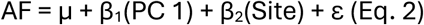

We report the *P*-values for this comparison alongside the *P*-values from 100 permutations, in which the habitat labels were randomized. SNPs were considered significant if their *P*-values in the real data outperformed those of at least 95% of the permutations. Next, we calculated the pairwise *F*_ST_ between the two populations and identified the top 10% most differentiated SNPs. Finally, we curated a list of candidate SNPs that were significant in both the GLM analysis and the top 10% of the *F*_ST_ distribution. We annotated all candidate SNPs using a gene feature file generated using AUGUSTUS *ab initio* gene prediction [(Stanke et al., 2008); model trained in *D. melanogaster*], SNPeff (REF: http://dx.doi.org/10.4161/fly.19695), as well as NCBI blastp searches to the *D. melanogaster* genome. We tested the set for enrichment of GO biological process terms with the Gene Ontology Resource online PANTHER tool(Mi et al., 2013); https://www.geneontology.org/). Lastly, as an additional layer of curation, for all genes identified as associated with outlier SNPs, we mined data from the FlyAtlas2 Anatomical Expression Data (Krause et al., 2022) to determine the cell and/or tissue types in which the gene was predominantly expressed.

### Metagenomic Analysis of Fly Microbiomes

High-throughput metagenomic sequencing and data analysis were conducted on bacterial and fungal communities from four sample types: wild-caught *D. basisetae*, baits used to capture flies, cultured slurry inocula, and lab-treated flies (**Fig. 1**). The full methods and primers can be found in Medeiros *et al*. (2024). Wild flies were collected from Tom’s Trail and Ola’a locations (see methods above) and transferred into individual microfuge tubes containing 95% EtOH in the field and then transported to the laboratory and stored at -80 _o_C until processing (**Fig. 1**). For DNA extraction, the flies were surface sterilized (2 X 95% EtOH) and rinsed (2 X MilliQ H2O) prior to homogenization. The homogenate was lysed and extracted using the MagAttract PowerSoil DNA EP Kit (Qiagen). Samples from banana and mushroom baits at the two sites were collected at the same time and processed in the same manner as the flies for microbiome analyses. Slurry inocula were generated by housing individual wild flies for 8 h and then washing the vials in sterile PBS, pooling fecal washes from ten flies per site, culturing aliquots on MRS (bacteria) or YPD (fungi) media (48 h at 18 °C then 18 h at 24 °C), and collecting colonies into a PBS:glycerol (1:1) solution before freezing at –80 °C; DNA was then extracted from these pooled slurries. Lab-treated flies (n = 6 per treatment) were snap-frozen at –80 °C immediately after the final slurry inoculation and temperature acclimation, surface sterilized, homogenized, and DNA-extracted identically to wild flies (**Fig. 1**; see below).

For each of the microbiome samples, bacteria were characterized by PCR amplification of the 16S rRNA gene and fungi by amplifying the internal transcribed spacer (ITS). Bacterial and fungal diversity were assessed based on PCR amplification of the 16S rRNA gene using, respectively, primers to the V3-V4 region (515F: GTGYCAGCMGCCGCGGTAA; 806R: GGACTACNVGGGTWTCTAAT) (Parada et al., 2016) and amplification of the internal transcribed spacer was performed (ITS1f: CTTGGTCATTTAGAGGAAGTAA; ITS2: GCTGCGTTCTTCATCGATGC) (White et al., 1990)). The primers contain a 12-base pair Golay-indexed code for demultiplexing. The PCRs were performed with the KAPA3G Plant kit (Sigma Aldrich, MO, USA) using the following conditions: 95 °C for 3 min, followed by 35 cycles of 95 °C for 20 seconds, 50 °C for 15 seconds, 72 °C for 30 seconds, and a final extension for 72 °C for 3 min. The PCR products were cleaned and normalized with the Just-a-plate kit (Charm Biotech, MO, USA). High throughput sequencing (HTS) was performed with Illumina MiSeq and 250 bp paired-end kits (Illumina, Inc., CA, USA). Post-processing of HTS data (read filtering, denoising, and merging) was performed using the “MetaFlow|mics’’ microbial 16S pipeline for bacteria and the fungal ITS pipeline for fungi (Arisdakessian et al., 2020). After post-processing, we chose a 97% sequence similarity cutoff for determination of operational taxonomic units (OTUs). Alpha and beta diversity analyses as well as relative quantities of microbial genera were analyzed in R using the phyloseq package (McMurdie & Holmes, 2013). Alpha diversity was measured using Chao1 and Shannon diversity metrics and a Wilcoxon statistical test. Beta diversity was calculated using non-metric multi-dimensional scaling (binary Jaccard distances) and an analysis of similarities (ANOSIM) test. Relative abundance charts were constructed after grouping flies from the same location, with univariate multiple testing and an *F*-test to test for significant differences in microbial taxa (phyloseq). Rarefaction curves for these samples were computed using the rarecurve command in phyloseq and using non-subsampled data, for the bacterial and fungal taxa from Tom’s Trail and Ola’a locations. In addition, a separate analysis was performed with four random samples from Tom’s Trail along with the original four samples from Ola’a (see **Fig. S4**).

### Drosophila husbandry

In the laboratory, the wild-caught flies from both locations were transported to UNLV and immediately placed in separate one-gallon breeding jars kept in a climate-controlled walk-in chamber and maintained at 18 °C and 60-70% RH, the standard rearing temperature for picture-winged *Drosophila*, with a 12 hr/ 12 hr day/ night cycle (**Fig. 1**;(J. Eldon et al., 2019; Uy et al., 2015). Each jar contained a layer of sand to provide moisture (RH in the jars was between 70-80%) and vials containing a standard adult Hawaiian *Drosophila* agar food medium [3 g agar, 6 g powder mix (equal portions of organic wheat germ, ground Special K® brand cereal, organic soy protein powder), 50 g Gerber® brand banana baby food, 90 ml ddH_2_O, 750 µl propionic acid, 750 µl ethanol] with a tissue soaked in a tea made from the bark and leaves of the larval host plant as is commonly required for raising Hawaiian picture wing *Drosophila* (Price & Boake, 1995). These adult food vials were replaced every 7 days, and those that contained larvae were fed Hawaiian *Drosophila* larvae food (2 g agar, 75 mL ddH_2_O, 20 g organic cornmeal, 2.2 g organic roasted soybean meal, 2.5 g organic nutritional brewer’s yeast, 5 mL Karo® brand corn syrup, 1.25 mL unsulfured molasses, 100 ml ddH_2_O) (Price & Boake, 1995) for approximately 3 weeks until 3^rd^ instar larvae began to wander out of the food. The larvae vials were then transferred to 1-gallon pupation jars that contained a one-inch layer of moist white gravel in which the third instar larvae were allowed to move out of the food vials and into the gravel to pupate. Adults that emerged within these pupation jars, after approximately 20 days as pupae, were transferred to new breeding jars or used in experiments. Laboratory populations were maintained in this controlled environment at a concentration of approximately 80-100 adults per jar with 10-15 adult jars for each population at any given time for 3 generations prior to experiments. The wild-caught founders were placed in 95% EtOH within 12 hr of death and stored at −20 °C for subsequent DNA extraction and analysis. For experimental assays, *D. basisetae* were removed from rearing jars within one week of eclosion, sexed, and placed into gallon-sized glass jars lined along the bottom with moist sand and provided with adult food vials. Hawaiian picture-winged *Drosophila* become sexually mature at approximately three-weeks of age (Craddock & Boake, 1992).

### Feces collection and bacteria and fungi culturing

To collect and culture bacteria and fungi from the feces of *Drosophila*, wild collected *D. basisetae* from each location were transported to the laboratory at UNLV and individually housed in sterile glass vials for 8 hours. After this period, the flies were removed, and the feces remaining in the vials were washed with sterile phosphate-buffered saline (PBS). The resulting feces-PBS solution was transferred to 1.5 mL microfuge tubes and stored in 500 µL of 1:1 ratio of PBS & glycerol. For microbial culture, aliquots of the feces solution were plated onto MRS (de Man, Rogosa, and Sharpe media: MilliporeSigma) plates to culture bacteria and YPD (Yeast Peptone Dextrose media: RPI Research Products) plates to culture fungi. The plates were incubated at 18 °C for 48 hours, followed by an additional 18 hours at 24 °C to promote microbial growth. All bacterial and fungal colonies were collected from the plates, and the cultures were pooled from ten individual flies from each location to generate combined samples for further analysis. These pooled bacterial and fungal slurries for both Tom’s Trail and Ola’a locations were used in microbe inoculation experimental procedures.

### Microbiome Inoculation and Temperature Acclimation Survival

For microbiome inoculation experiments, virgin female and male flies were first separated by sex within 7 days of eclosion and placed in adult food vials for three days at 18 °C and then subsequently placed in vials with an antimicrobial diet for 7 days and housed in an incubator at 18°C (**Fig. 1**). The antimicrobial diet consisted of a standard diet that was supplemented with 1.25 mM captan (N-trichloromethylmercapto-4-cyclohexene-l,2-dicarboximide; Sigma-Aldrich), an antifungal chemical, suspended in coconut oil (300 μL/ L standard media; Kirkland brand), and three anti-bacterial compounds: 200 mg/ mL of ampicillin and kanamycin (VWR Life Science; both dissolved in water), 50 mg/ mL of tetracycline (EMD Millipore Corp.; dissolved in 70% ethanol), and 300 mg/ mL erythromycin (Acros Organics; dissolved in 100% ethanol). This antimicrobial diet has been shown to dramatically reduce both fungi and bacteria in Hawaiian picture-winged *Drosophila* (Medeiros et al., 2024).

After the antimicrobial treatment, male and female *D. basisetae* from Tom’s Trail and Ola’a populations were randomly placed (1 or 2 flies per vial) in sterilized glass vials containing a standard diet supplemented with 12.5 µL of a microbiome slurry from either Tom’s Trail or Ola’a pipetted onto the surface of the food (see above). The vials, along with the flies, were randomly placed in either an 18 °C or 24 °C Percival incubator for 15 days; RH was 60-70% for both incubators. The lower or control temperature of 18 °C is a typical standard temperature for raising Hawaiian picture wing *Drosophila* (Price & Boake, 1995). The high temperature of 24 °C was chosen as a stressful temperature has been shown in previous studies to cause differences in survival and reproductive traits between Hawaiian picture wing *Drosophila* species and between populations collected from similar elevations as this current study (J. Eldon et al., 2019; Uy et al., 2015). During these 15 days, every three to four days (alternating between three and four days), the flies were transferred to fresh sterile vials containing the same food and microbiome slurry. Every two to three days, the survival of the flies was recorded.

### Basal Movement and Critical Thermal Maximum

To determine the basal movement at control temperature and critical thermal maximum (CTmax) of *D. basisetae*, flies that had been acclimated to either 18 °C or 24 °C and the two microbiome inoculation treatments, were next placed in 18 °C for three days. After three days of acclimation to the control temperature the flies were placed individually into the wells of a tissue culture plate (2 cm diameter), with each plate housing 12 flies. A total of eight plates at a time, accommodating 96 flies, were placed inside a Darwin KB011 chamber (Darwin Chambers, St. Louis, MO, USA) equipped with two video cameras mounted above the tissue culture plates. The video cameras (Weewooday Camera 1080P Webcam 5MP OV5647 Sensor Day and Night Vision IR-Cut Video Camera) were each connected via a video cable to a Raspberry Pi 4 (CanaKit) computer and video monitor outside the chamber. The temperature and humidity experienced by the flies during the procedure was recorded by an Elitech Digital Temperature and Humidity Data Logger GSP-6 with the temperature and humidity probes in a tissue culture plate inside the Darwin Chamber. After the flies were placed in the tissue culture wells and transferred to the Darwin chamber, the Raspberry Pi video recording and Elitech data logger was initiated. The Darwin chamber was programmed to initiate the temperature at 18 °C for 15 minutes to allow the flies to acclimate for 5 minutes, then the basal movement of the flies was recorded at the control temperature for 10 minutes. The temperature in the chamber was gradually increased at a rate of 0.5 °C per minute for 40 minutes and then slowed to a ramp of 0.25 °C per minute for the next 60 minutes. The basal movement and CTmax for each fly was determined by analyzing video recordings and the temperature data with EthoVision XT software (Noldus) to identify the movement for 10 min. at the beginning of each trial, and when movement of each fly had ceased, indicating the CTmax for each individual fly. After the CTmax procedure, each fly was transferred to a 1.5 μL microfuge tube with PBS and stored in 4 °C refrigerators for subsequent measurements.

### Mature egg counts and Accessory gland size measurements

The flies that were stored in PBS in a 4 °C refrigerator after CTmax measurements were dissected within 5 days. The number of mature eggs was manually counted in dissected females under 10x magnification using a Nikon SMZ25 dissecting microscope connected to a Windows computer with Nikon NIS Elements version 4.6 video software. The male accessory glands were also measured within 5 days after CTmax measurements. The males were dissected and the accessory glands were carefully positioned under the Nikon dissecting microscope and a picture was taken that included a micrometer for size standardization. The accessory gland images were saved and later measured with ImageJ software (Schneider et al., 2012). The size was measured by outlining the entire accessory gland image and the area of the accessory gland was calculated in the ImageJ software.

General Linear Models (GLM) in Minitab (Minitab® 21.4.3) were used to assess the significance of fly source population, temperature treatment, microbiome treatment, sex (for non-reproductive traits) and their interactions on the phenotypic measurements: acclimation survival, basal movement, CTmax, mature egg number, and accessory gland size. To obtain sufficient sample size, the microbiome and temperature acclimation experimental procedures and phenotypic measurements were conducted over five different times (blocks). For each response variables the block effects was included as a fixed effect to account for differences among the weeks of the measurements in the GLM analysis.

## RESULTS

### Population Genomic Analysis of Wild Flies

To assess the extent of genetic differentiation and potential signals of local adaptation between populations, we conducted population genomic analyses of wild *D. basisetae* flies collected from Tom’s Trail and Ola’a. We genotyped *D. basisetae* samples using two approaches. The GATK-haplotype method yielded 4,520,251 SNPs before the VCF filtering step. After applying our stringent VCF filters (see Materials and Methods), we obtained 375,730 high-quality genome-wide SNPs with coverage ≥4X. We used these data to survey population structure within and among our samples. We report the fixation index (*F*_ST_) as well as two multidimensional ordination methods, PCA (unsupervised clustering) and DAPC (supervised clustering). The latter is a method well-suited for detecting latent population structure (Jombart, 2008). Both the results from PCA and *F*_ST_ reveal evidence for weak population structure (*F*_ST_ = 0.0084; **Fig. 2A**). For example, both populations have a continuous ordination pattern in PC 1 (6.39% Variance Explained [VE]) with a weak clustering pattern whereby samples from Tom’s Trail are more closely clustered with each other (Euclidean mean distance [*d*] among Tom’s Trail samples = 359.90; sd*_d_* = 76.52) than those from Ola’a (*d* = 479.21; sd*_d_* = 25.45). These patterns are not driven by family structure in our samples (mean kinship coefficients in Ola’a = -0.0763, Tom’s = -0.0642).

**Fig 2.**
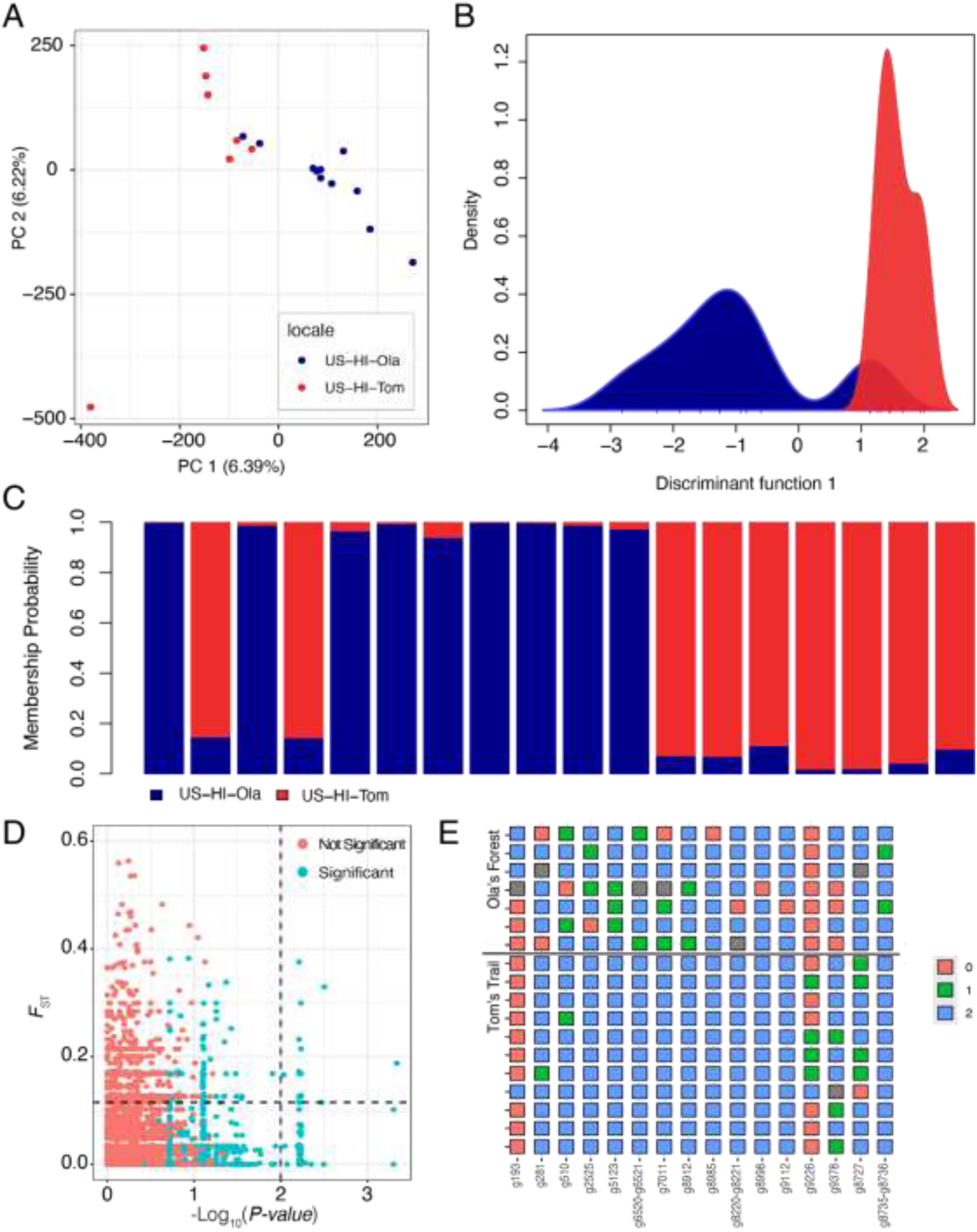
(A) Principal component analysis on SNPs derived from the GATK-haplotype pipeline. (B) Discriminant component analyses on the same data as A. (C) Membership probability analysis from the DAPC model. The color scheme is shared between A, B, and C with blue being Ola’a and red being Tom’s Trail. (D) Co-localization plot showing the pairwise FST between the two populations and the -Log_10_(P-value) of the GLM test. The red dots indicate SNPs that do not beat permutations. (E) Genotype plots for the 16 outlier SNPs identified in our analyses. The number and color indicate the number of alleles in the sample relative to the reference (0 = homozygous reference; 2 = homozygous alternative). Grey indicates missing data.

To further explore these patterns, we conducted a DAPC on the same data. Results from the DAPC are overall consistent with those of the PCA. Accordingly, the DAPC algorithm generated only one synthetic discriminant function that mirrors the patterns from PCA (**Fig. 2B).** Indeed, the sample projections from the DAPC are significantly correlated to those from PC 1 in the PCA (correlation [*r*] = -0.8707, *P* = 2.60x10-6). The membership posteriors derived from the DAPC model (**Fig. 2C**; i.e., the probability that a sample is classified as a member of a given site, contingent on its genetic makeup) shows evidence for gene flow and admixture between populations. For example, out of the 11 samples collected from Ola’a, only 7 showed membership posteriors of 99% or above to the Ola’a population. The other 4 samples show evidence of genetic admixture, with membership posteriors up to 50% for the Tom’s Trail population. Notably, samples collected at Tom’s Trail show a similar result, although membership posteriors for admixed samples are smaller than those at Ola’a (i.e., 1%-15%). This finding may suggest asymmetrical levels of gene flow among the sites.

Our second genotyping approach used “shallow-sequencing” likelihood-based methods to conduct population genetic analyses. As above, we capitalized on conservative filters to estimate genotype likelihoods from 220,261 SNPs. Despite the different analytical pipelines deployed, our results from the genotype-likelihood analyses are highly concordant with those from the haplotype-based pipeline. For example, we observe weak clustering patterns in PCA (**Fig. S1**) and very low levels of *F*_ST_ (0.039). Overall, these findings suggest that Ola’s Forest and Tom’s Trail are weakly structured populations that are connected by gene flow.

We implemented a battery of tests to explore potential targets of local adaptation. Given the properties of our data, we implemented stringent filters to assess significance in our GLM model and identify potential outlier SNPs. Overall, we identified 167 SNPs across the genome of *D. basisetae* with *P*-values in our GLM test that beat 95% of permutations. Of these, only 16 SNPs, across 18 genes, are also highly differentiated by *F*_ST_ (90^th^ quantile, *F*_ST_ > 0.114; **Fig. 2D-E**). These SNPs were largely predicted to have a modifier functional effect, with three intergenic SNPs, five located in introns, one located in a 5’ UTR, and one synonymous variant. Notably, these loci are not dispersed across the genome, since most are concentrated on scaffold JAWNLB010000174.1 (10 SNPs 58%), two SNPs are located in JAWNLB010000013.1, and the remainder are in unique scaffolds. Genes associated with the 16 SNPs are involved with a variety of functions in *D. melanogaster*, including immunity, reproduction, physiology, neuronal signaling and behavior (**Table S2**). There were no significantly enriched GO terms represented within the set.

### Microbiomes of Wild Flies

We next examined the structure of microbial communities associated with wild flies from both locations to assess how microbiome diversity and composition vary across populations and environments. High-throughput amplicon sequencing analysis revealed significant differences in the bacterial and fungal components of flies collected from Tom’s Trail and Ola’a, both in terms of taxonomic diversity and composition.

***Bacterial Communities***: Alpha diversity measures of bacterial taxa revealed significantly lower diversity at Ola’a compared to Tom’s Trail (Chao 1, *P* = 0.049), although Shannon diversity did not differ between the two sites (*P* = 0.72; **Fig. 3A**). Measurement of beta diversity of bacterial communities also showed significant differences between Tom’s Trail and Ola’a (ANOSIM using Jaccard’s distance, *P* = 0.047; **Fig. 3B**). The relative frequencies of the ten most common bacterial taxa trended differently between the two locations though the differences were not significant. *Gluconobacter* and *Pseudomonas* tended to be more abundant in Ola’a (31.4% and 25.1%, respectively) than in Tom’s Trail (18.7% and 4.2%, respectively). Other bacteria, such as *Leuconostoc* and *Gilliamella*, were relatively more abundant at Tom’s Trail (26.1% and 6.4%, respectively) than at Ola’a (11.2% and 0.7%, respectively) (**Table 2**, **Fig. 3C**).

**Fig 3.**
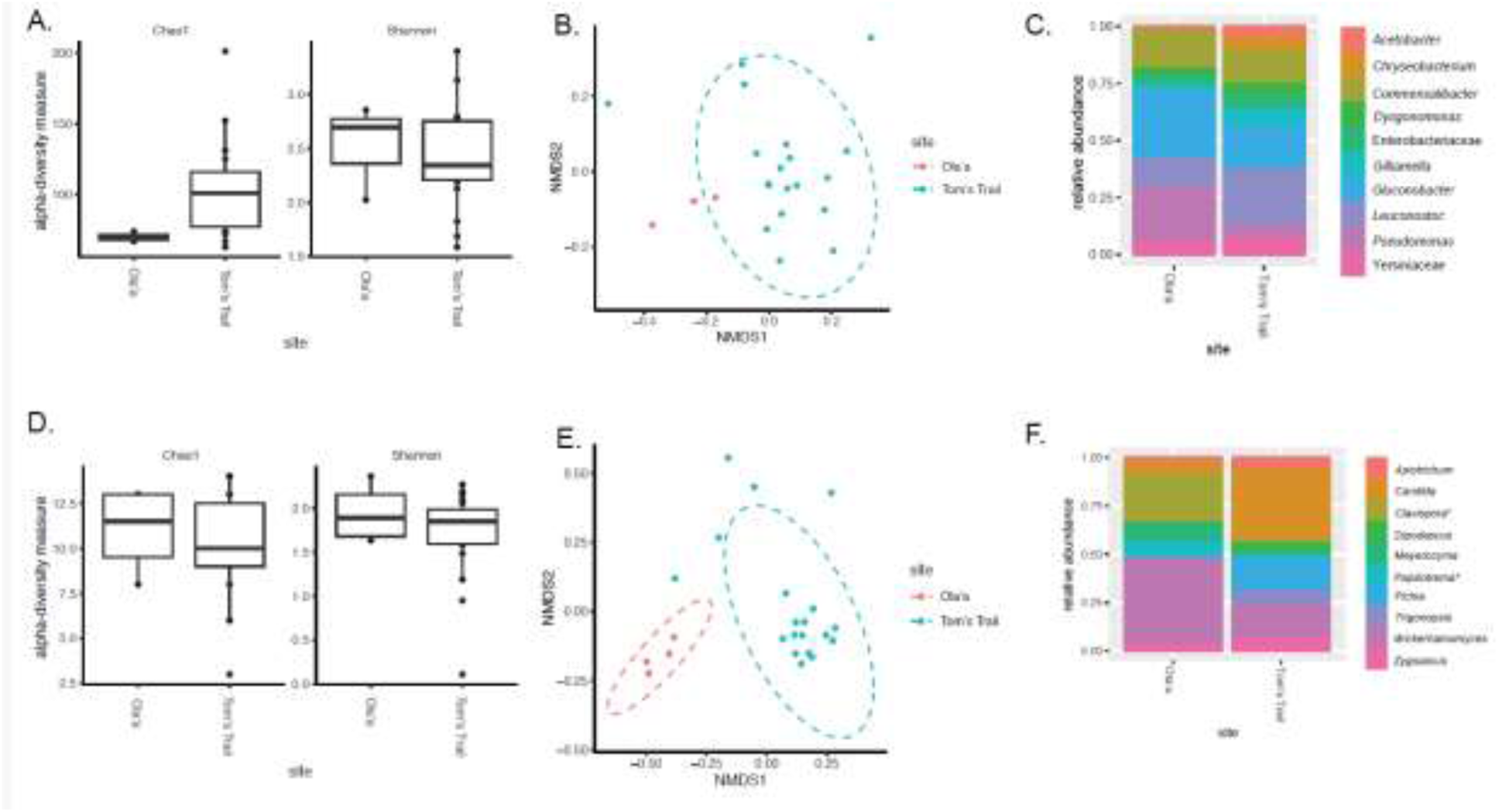
Microbiome of wild *D. basisetae* flies: Alpha and Beta Diversity measures of bacterial and fungal taxa from Tom’s Trail & Ola’a. (A) Taxonomic richness (based on Chao1) of bacterial taxa was significantly lower at Ola’a compared to Tom’s Trail (*P* = 0.049); Shannon diversity did not differ between sites (*P* = 0.72). (B) Beta Diversity of bacterial taxa (based on ANOSIM using Jaccard’s distance) differed significantly between sites (*P* = 0.047). (C) The relative frequencies of the ten most common bacterial taxa in both locations. Alpha and Beta Diversity measures of Fungi from Tom’s Trail & Olaʻa. No significant differences were detected. (D) Taxonomic richness (based on Chao1) of fungal taxa was not significantly different between sites (*P* = 0.77) nor was Shannon diversity (*P* = 0.41). (E) Beta Diversity of fungi (based on ANOSIM using Jaccard’s distance) was highly significantly different between sites (*P* = 0.002). (F) The relative frequencies of the ten most common fungal taxa in both locations; *: *Clavispora* (*P* = 0.001) and *Papiliotrema* (*P* = 0.02) differed significantly. N=19 for Tom’s Trail and n=4 for Olaʻa.

**Table 2.**
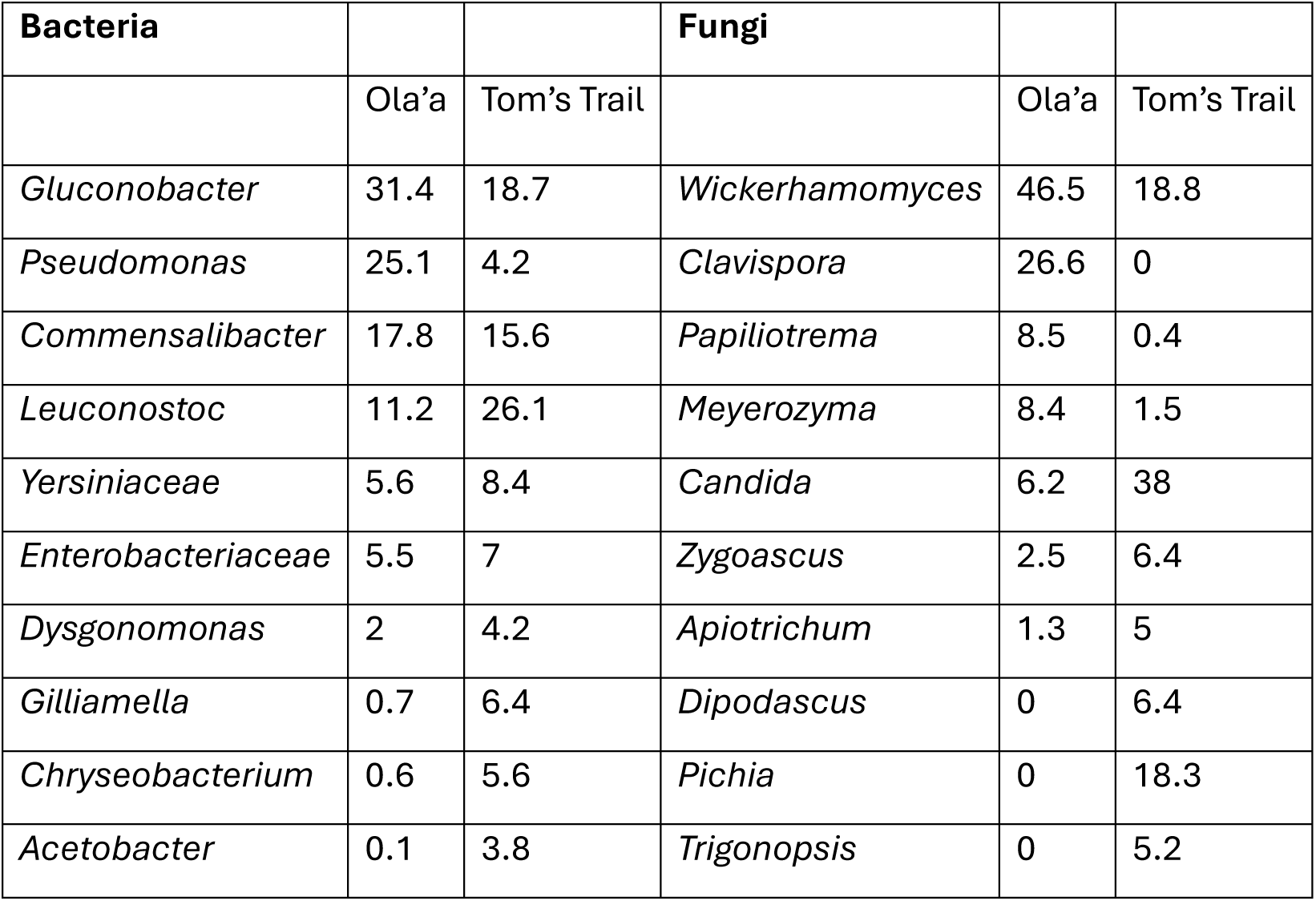
The relative frequencies of ten most common bacteria and fungi collected from *D. basisetae* flies.

| **Bacteria** |  |  | **Fungi** |  |  |
| --- | --- | --- | --- | --- | --- |
|  | Ola'a | Tom's Trail |  | Ola'a | Tom's Trail |
| *Gluconobacter* | 31.4 | 18.7 | *Wickerhamomyces* | 46.5 | 18.8 |
| *Pseudomonas* | 25.1 | 4.2 | *Clavispora* | 26.6 | 0 |
| *Commensalibacter* | 17.8 | 15.6 | *Papiliotrema* | 8.5 | 0.4 |
| *Leuconostoc* | 11.2 | 26.1 | *Meyerozyma* | 8.4 | 1.5 |
| *Yersiniaceae* | 5.6 | 8.4 | *Candida* | 6.2 | 38 |
| *Enterobacteriaceae* | 5.5 | 7 | *Zygoascus* | 2.5 | 6.4 |
| *Dysgonomonas* | 2 | 4.2 | *Apiotrichum* | 1.3 | 5 |
| *Gilliamella* | 0.7 | 6.4 | *Dipodascus* | 0 | 6.4 |
| *Chryseobacterium* | 0.6 | 5.6 | *Pichia* | 0 | 18.3 |
| *Acetobacter* | 0.1 | 3.8 | *Trigonopsis* | 0 | 5.2 |

***Fungal Communities*:** Alpha diversity measures of fungal taxa did not differ significantly between Tom’s Trail and Ola’a (Chao1, *P* = 0.77; Shannon diversity, *P* = 0.41; **Fig. 3D**). However, Beta diversity of fungal taxa was highly significantly different between the two locations (*P* = 0.002; **Fig. 3E**). The relative frequencies of the ten most common fungal taxa varied between Tom’s Trail and Ola’a. *Wickerhamomyces* and *Clavispora* were the most abundant fungal taxon at Ola’a (46.5% and 26.6%, respectively) compared to Tom’s Trail (18.8%, 0%), while *Candida* was more abundant at Tom’s Trail (38%) than Ola’a (6.2%). *Pichia* was relatively common at Tom’s Trail but were not detected from the Ola’a community. Additionally, *Dipodascus* and *Trigonopsis* were present at Tom’s Trail but were undetectable from the fungal community at Ola’a (**Table 2**, **Fig. 3F**). However, only *Clavispora* and *Papiliotrema* differed significantly. Rarefaction curves for the bacterial and fungal taxa from these samples show the same general pattern with greater number of bacterial and fungal taxa at Tom’s Trail location compared to Ola’a and indicating that the sequencing analyses were sufficient to detect most taxa in the samples (**Fig. 3S**).

A comparison between microbes in flies and baits revealed notable distinctions (**Table S3**). While taxa such as *Leuconostoc*, *Gluconobacter*, and *Pseudomonas* occurred in both, specific OTUs of some of these shared genera differed between flies and baits. Additionally, certain bacterial taxa (e.g., *Halomonas*, *Comamonas*, *Sphingobacterium*, and specific OTUs of *Stenotrophomonas*) and fungal taxa (e.g., *Limtongozyma*, *Geotrichum*, *Papiliotrema*) were exclusive to flies. Conversely, other bacterial taxa (e.g., specific OTUs of *Weissella*, *Lactococcus*, *Brevundimonas*) and fungal taxa (e.g., specific OTUs of *Penicillium*, *Saccharomycopsis*, *Wickerhamomyces*) appeared only in baits, supporting the idea that fly microbes are acquired from environmental sources other than bait.

### Microbe Composition of Slurries and Treated Flies

To assess how microbiomes may influence thermal tolerance and reproduction, we next characterized microbial communities in the slurries used to inoculate *D. basisetae* and then examined the microbiome of laboratory flies after the antimicrobial and inoculation treatment.

***Slurry Microbes Used for Inoculation*:** The slurries used in the inoculation experimental procedures were made by culturing bacteria and fungi from feces of wild flies. These slurry communities differed considerably between locations (**Fig. S2**). The Olaʻa slurry was dominated by *Gluconobacter* (81%) and *Bacillus* (17.3%) among bacterial taxa, while the Tom’s Trail slurry was more diverse and dominated by *Leuconostoc* (40%) and *Psychrobacter* (40%), while *Gluconobacter* was much less abundant (11%) and minor contributions from *Morganella* (3%), *Acetobacter* (3%), and *Lactobacillus* (3%) (**Fig. S2A**). Fungal communities in the slurries also differed: Olaʻa slurries were composed almost entirely of *Pichia* (97%) with a small proportion of *Penicillium* (3%), while Tom’s Trail slurries were dominated by *Dipodascus* (79%) with *Pichia* making up the remainder (21%) (**Fig. S2B**). These cultured slurry profiles were substantially different from the wild fly microbiomes (**Fig. 3**), likely due to differential growth on MRS and YPD media.

***Microbiomes of Treated Laboratory Flies*:** We then analyzed the microbiomes of lab-reared flies following microbial inoculation and thermal acclimation to assess whether microbiome composition remained distinct post-inoculation (**Fig. 4**). Bacterial alpha diversity (Chao1 and Shannon indices) did not differ significantly across treatments (**Fig. 4A**), nor did beta diversity (Jaccard-based ANOSIM, **Fig. 4B**). The relative abundance of the six major bacterial taxa remained similar between treatment groups (**Fig. 4C**). In contrast, fungal communities showed more pronounced differences. Alpha diversity of fungi did not differ significantly across treatments (**Fig. 4D**), but beta diversity differed significantly between groups (ANOSIM, *P* < 0.001; **Fig. 4E**). Notably, all pairwise comparisons of fungal communities were significantly different (*P* = 0.01) except when flies received slurries from their site of origin. This pattern suggests that fungal communities were more responsive to the interaction of host population and microbial source. Differences in the relative abundance of specific fungal taxa were also detected: *Pichia* (*P* = 0.0002) and *Geotrichum* (*P* = 0.0335) varied significantly between treatments (**Fig. 4F**).

**Fig 4.**
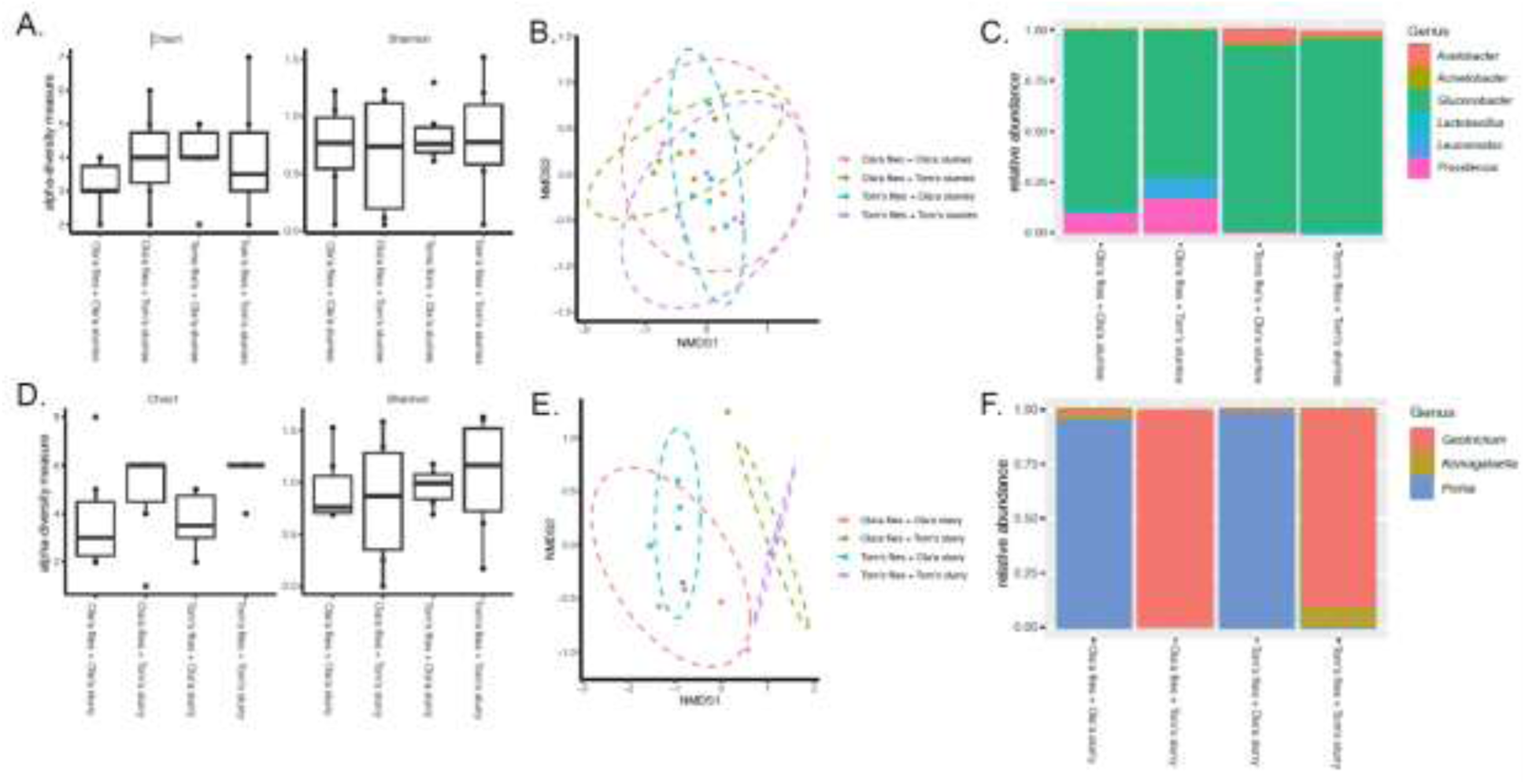
Microbiome of lab *D. basisetae* flies after receiving fecal slurries from either Tom’s Trail or Olaʻa, n=6 for all treatments. A-C report 16S rRNA analyses and D-F report ITS analyses. (A) Taxonomic richness of bacterial taxa (Chao1 and Shannon diversity) did not differ significantly between treatments. (B) Beta Diversity of bacterial taxa (based on ANOSIM using Jaccard’s distance) did not differ significantly between treatments. (C) The relative frequencies of the six bacterial taxa did not differ between treatments. (D) Taxonomic richness of fungal taxa (Chao1 and Shannon diversity) did not differ significantly between treatments. (E) Beta Diversity of fungi (based on ANOSIM using Jaccard’s distance) was different between treatments (*P*<0.001) with all pairwise comparisons differing significantly (*P*=0.01) except those receiving the same fecal slurries (Olaʻa flies+Olaʻa slurry vs. Tom’s flies+Olaʻa slurry and Olaʻa flies+Tom’s slurry vs. Tom’s flies+Tom’s slurry). (F) The relative frequencies of the three fungal taxa present varied between treatments in two cases: *Pichia* (*P*=0.0002) and *Geotrichum* (*P*=0.0335).

Together, these findings demonstrate that while bacterial communities in treated flies remained relatively similar across inoculation conditions, fungal communities were substantially different between microbial source and environmental context. These microbiome differences may help explain the phenotypic variation observed in thermal tolerance and reproductive traits across experimental treatments.

### Phenotypic Analyses of Experimental *D. basisetae*

To evaluate how microbial composition and environmental temperature interact with host traits, we analyzed survival, behavior, and reproductive traits in the *D. basisetae* laboratory populations initiated from wild flies collected from Tom’s trail and Ola’a sites and exposed to a combination of microbial inoculation of slurries originating from Tom’s trail or Ola’a fly populations, and temperature acclimation treatments of 18 °C or 24 °C. Females and males from these laboratory populations showed significant microbiome inoculation and temperature acclimation treatment effects. The flies from both locations that were exposed to the Tom’s Trail microbes, compared to Ola’a microbes, had significantly higher survival during the 15 days of the acclimation period (*F_1,242_* = 4.68, *P* = 0.031) and a trend for greater survival at 18 °C compared to 24 °C (**Fig. 5A**; *F_1,242_* = 3.25, *P* = 0.079): there was no main effect of sex (*F_1,242_* = 0.26, *P* = 0.611) or fly population (*F_1,242_* = 2.53, *P* = 0.113). For fly basal locomotion, there was no main effect of sex (*F_1,50_* = 0.30, *P* = 0.584), fly population (*F_1,50_* = 0.51, *P* = 0.478) or temperature (*F_1,50_* = 0.08, *P* = 0.777) but a significant interaction between fly population and microbe treatment (*F_1,50_* = 6.75, *P* = 0.012). Flies exposed to the microbes from their original location moved more than flies exposed to microbes from the other location (**Fig. 5B)**. In the case of CTmax, there were main no effects of sex (*F_1,50_* = 1.63, *P* = 0.208), fly population (*F_1,50_* = 1.54, *P* = 0.221), or temperature (*F_1,50_* = 1.15, *P* = 0.289) but a significant three-way interaction between fly population, temperature treatment, and microbiome treatment (*F_1,50_* = 4.21, *P* = 0.045), with flies from Tom’s Trail location showing the greatest interaction of temperature and microbiome treatment effect (**Fig. 5C**).

**Fig 5.**
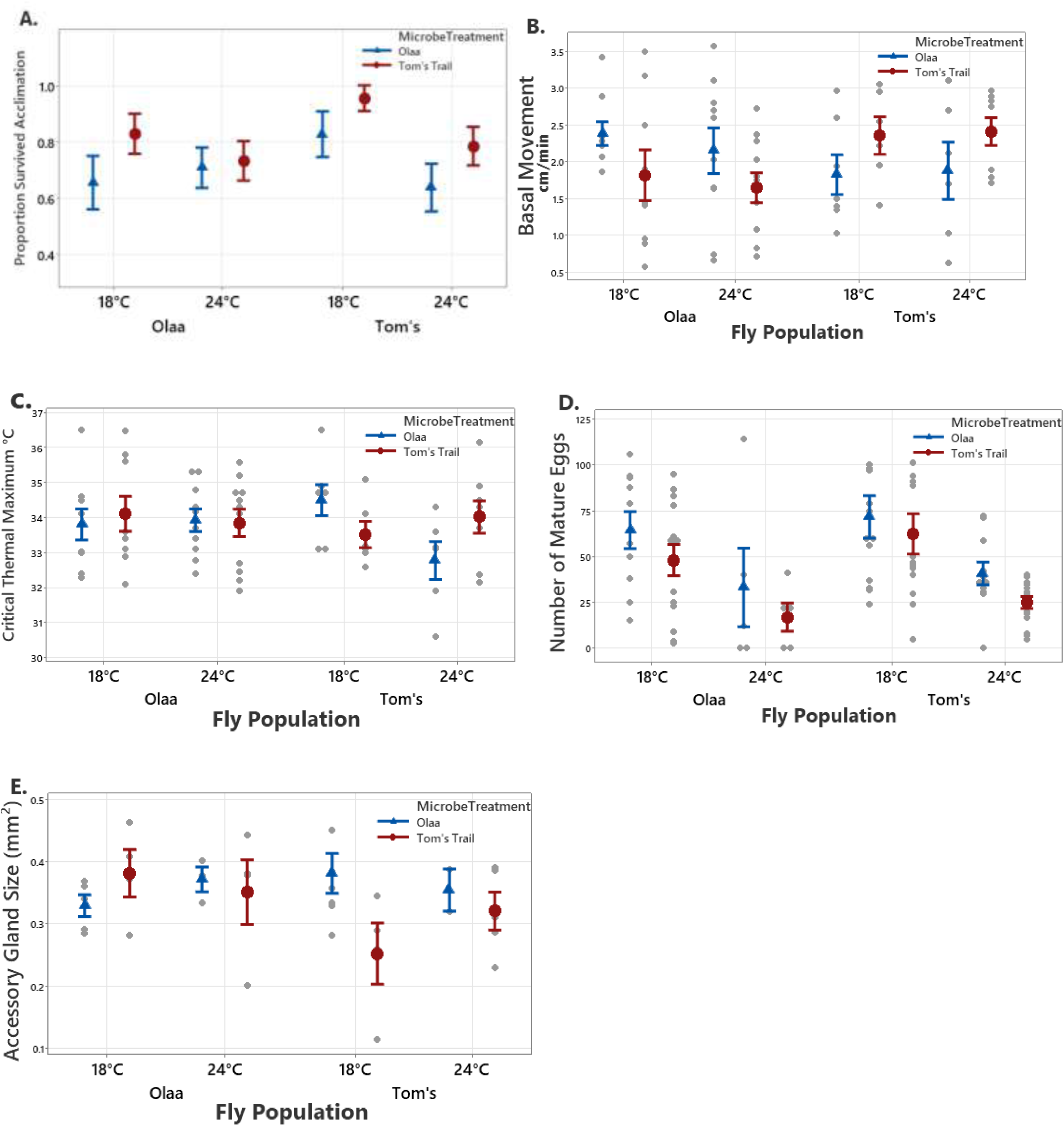
General Linear Model (GLM) of phenotypic traits measured on *D. basisetae* laboratory treatment groups that originated from Ola’a and Tom’s Trail populations. The means and standard errors of the traits measured on flies from these laboratory populations are shown that were given microbe treatment from Ola’a (blue triangle) or Tom’s Trail (red circle) cultured slurries and acclimated at 18 °C or 24 °C. (A) The proportion of individuals that survived 15 days of acclimation, (B) the basal movement of flies for 10 minutes at 18 °C, (C) the critical thermal maximum (CTmax) measured during a temperature ramping protocol, (D) the number of mature eggs of females at approximately 25 days old, (E) the size of the male accessory gland measured at 25 days old. The small grey circles indicate the individual values basal movement, critical thermal maximum, number of mature eggs and accessory gland size. The individual values for proportion survived acclimation were either 0 or 1 and are not shown.

For the number of mature eggs, there was no main effect of fly population (*F*_1,84_ = 0.72, *P* = 0.398) but there was a highly significant main effect of temperature with more eggs produced in flies housed at 18 °C compared to 24 °C (*F*_1,84_ = 19.92, *P* = 0.001), as well as a main effect of microbe treatment with more mature eggs produced when flies were given Ola’a microbes compared to Tom’s Trail microbes (**Fig. 5D**; *F*_1,84_ = 4.07, *P* = 0.047) and none of the interaction effects were significant (P > 0.30). For male accessory gland size there were no main effects of fly population (*F*_1,33_ = 0.35, *P-value* = 0.558), microbe treatment (*F*_1,33_ = 1.42, *P* = 0.242) or temperature (*F*_1,33_ = 0.65, *P* = 0.427) but there was a significant three-way interaction of fly population, temperature treatment, and microbiome treatment (*F*_1,33_ = 4.43, *P* = 0.043) with flies from Tom’s Trail showing the greatest interaction effect (**Fig. 5E)**. These results underscore the importance of both temperature and microbe treatment in shaping key behavioral, reproductive, and physiological traits in *D. basisetae* in different populations.

## Discussion

This study provides insights into how microbiomes may contribute to local adaptation and phenotypic plasticity in *Drosophila basisetae*. By integrating microbiome inoculation, temperature acclimation, and population genomic analyses, we explored how microbial communities sourced from distinct habitats interact with host species population genomic divergence at key adaptive loci to influence host physiology. Although previous research has established that microbiomes are critical for insect health, metabolic efficiency, and reproductive fitness (Broderick & Lemaitre, 2012; Delbare et al., 2020; Douglas, 2015), our findings suggest that microbiome composition and its interaction with environmental factors could play a role in shaping host survival and reproductive traits during thermal stress (Bost et al., 2018; McMullen et al., 2020).

### Population Genomic Analysis

Population genomic analyses revealed that although there is relatively low overall genetic differentiation between the populations from Tom’s Trail and Ola’a, a number of SNPs showed significantly higher population divergence than expected, suggestive of population-specific adaptation that may be associated with their divergent environments and microbiome communities. Outlier SNPs were primarily located in gene regulatory regions, with potential impacts on a variety of functions that may be relevant for host-microbiome interactions. Two outlier SNPs appear directly related to host immunity. *SkpA* is part of the *Imd* pathway involved in regulation of the anti-bacterial peptide Diptericin; loss-of-function of this gene leads to constitutive expression of Diptericin, preventing down-regulation following recovery from infection (Aparicio et al., 2013; Khush et al., 2002). The protein yin is a phagosome-associated transporter involved in bacterial ligand sensing and transport (Charrière et al., 2010). The remaining genes belonged to several functional categories tied to the phenotypes showing interaction effects in this study, including heat tolerance (*unc-45*), reproduction (*APC7, pigs*), and neuronal functions related to behavior and memory (*pHCl-1, CARPA, dar1, klg, tsf1, Antdh, Tsp5D, retm, drp20*). Notably, *drp20*, and *drp* proteins more generally, have been linked to microbiome-mediated nutritional impacts on body size in *D. melanogaster* (Dobson et al., 2015), and are evolutionarily responsive to microbiome composition (Dobson et al., 2015; Rudman et al., 2019). It should be noted that our statistical power to detect local adaptation was limited by sample size, resulting in strict thresholds that likely excluded additional biologically relevant SNPs. However, despite this limitation, these candidate loci suggest localized adaptation that may interact with gut-microbe composition to alter host reproduction and survival abilities (Agler et al., 2016; Amend et al., 2022; J. Eldon et al., 2019; Uy et al., 2015). These findings are consistent with those of Eldon et al. (2019), who showed that adaptive divergence can occur over short distances and subtle climatic gradients in Hawaiian *Drosophila*. In that study, *D. sproati* populations, only 7 km apart and 365 m difference in elevation, exhibited strikingly different physiological, behavioral, and gene expression responses to a similar mild heat exposure despite low neutral genetic differentiation, suggesting that fine-scale environmental heterogeneity may drive localized adaptation, potentially mirrored in the microbiome-mediated responses we observed in *D. basisetae*.

### Microbial Community Composition and Functional Roles in Thermal Adaptation

Distinct differences in microbial community composition between *D basisetae* populations from Tom’s Trail and Olaʻa suggest that local environmental conditions influence the structure of both bacterial and fungal assemblages. Wild flies from Tom’s Trail harbored a more diverse bacterial community, with *Leuconostoc, Gluconobacter* and *Commensalibacter* among the dominant taxa, whereas flies from Olaʻa had a less diverse microbiota enriched primarily in *Gluconobacter* and *Pseudomonas* (Adair et al., 2018; Armstrong et al., 2022). Fungal communities also differed significantly between sites: *Wickerhamomyces* and *Clavispora* were abundant in Olaʻa flies, while Tom’s Trail flies showed greater diversity, including *Candida*, *Pichia,* and *Wickerhamomyces*. These compositional differences may be influenced by environmental factors such as elevation, rainfall, and habitat-specific microbial reservoirs, consistent with findings in other insect systems (Houwenhuyse et al., 2021; Martino et al., 2017; Medeiros *et al*., 2025).

Comparisons between microbes found in flies and those present in bait substrates revealed partial overlap, shared bacterial genera such as *Leuconostoc*, *Gluconobacter*, and *Pseudomonas* were detected in both, but several bacterial taxa (e.g., *Halomonas*, *Comamonas*, *Sphingobacterium*) and fungal taxa (e.g., *Geotrichum*, *Papiliotrema*) were exclusive to flies. These findings support the idea that *D. basisetae* may selectively acquire specific microbes from the environment, potentially influenced by host behavior or gut filtering mechanisms (Acevedo et al., 2021; Corby-Harris et al., 2007). Although the cultured slurries used for microbiome inoculation represented a simplified subset of the wild microbial communities, likely due to media selectivity, they retained differences. Slurries from Olaʻa were dominated by *Gluconobacter* and *Bacillus* bacteria taxa and *Pichia* fungal taxa, while those from Tom’s Trail featured *Psychrobacter*, *Leconostoc* and *Gluconobacter* bacteria taxa and the fungal taxa *Dipodascus* with less abundance of *Pichia*. When administered to lab-reared flies, bacterial communities remained relatively similar across treatments, but fungal communities showed marked differences depending on the microbial source. In particular, flies inoculated with Olaʻa slurries were dominated by *Pichia* fungal taxa while those from Tom’s Trail were dominated by *Geotrichum* fungal taxa, suggesting an important role of fungal taxa in mediating host-microbe interactions (Houwenhuyse et al., 2021; Medeiros et al., 2024; Walters et al., 2020). Future research should clarify the acquisition pathways and investigate the biogeographic distribution of fly microbiomes across the diverse Hawaiian environments (Acevedo et al., 2021; Medeiros et al., 2025).

Functionally, the observed microbial differences were associated with divergence in thermal tolerance and reproductive traits, suggesting that gut microbiota may influence key phenotypic responses critical for survival and reproduction in divergent environments. Both populations of *D. basisetae* exhibited higher survival rates during the acclimation period when exposed to microbes derived from Tom’s Trail flies at both the control temperature of 18 °C and the high stressful temperature of 24 °C that has been shown to be stressful for Hawaiian picture wing *Drosophila* species from similar elevations as this current study (Eldon et al., 2019; Uy et al., 2015). This enhanced survival may be related to the metabolic support provided by specific microbial taxa (Broderick & Lemaitre, 2012; Wong et al., 2013). In addition, Tom’s Trail flies exhibited greater CTmax when exposed to Tom’s Trail microbes when acclimated to the higher temperature, but greater CTmax when exposed to Ola’a microbes when acclimated to the lower control temperature. Ola’a flies showed no significant differences in CTmax for either microbe or temperature acclimation treatment. These findings align with prior research indicating that microbes modulate key metabolic pathways involved in stress tolerance and energy management (Martino et al., 2017; McMullen et al., 2020; Moghadam et al., 2018). These results also complement those of Uy et al. (2015), who found that similar heat stress in Hawaiian picture-winged *Drosophila* produced changes in gene expression, including upregulation of heat shock proteins. While Uy et al. (2015) focused on direct thermal responses, our findings suggest that microbiomes may act upstream or in parallel with these pathways to shape host plasticity in response to temperature.

Reproductive traits also showed differential responses depending on the microbe and temperature treatment, with flies from both locations producing more eggs at the cooler control temperature. Interestingly, flies from both populations also produced more eggs when exposed to the microbes from Ola’a compared to Tom’s Trail microbes, suggesting that *Pichia*-rich communities may promote reproductive investment. Male accessory gland size, a trait associated with male reproductive success, exhibited complex interactions between population origin, microbe treatment, and acclimation temperature, particularly in Tom’s Trail flies. These complex patterns suggest that different microbial communities may cause adaptive shifts in life-history strategies, possibly promoting either survival or reproductive investment based on microbiome composition and environmental context (Adair et al., 2018; Houwenhuyse et al., 2021; Téfit et al., 2023).

### Microbiome-Mediated Plasticity and Conservation Implications

Overall, our findings suggest that phenotypic plasticity associated with changes in the microbiome may enable *D. basisetae* populations to balance survival and reproduction in response to local environmental pressures (Kolodny & Schulenburg, 2020), possibly allowing rapid shifts in traits like thermal tolerance without significant genetic change (Armstrong et al., 2022; Petersen et al., 2023; Sommer & Newell, 2019). At the same time, adaptive genetic differences, also documented in other Hawaiian *Drosophila,* may contribute to these life-history strategies (Eldon et al., 2019; Price et al., 2014), underscoring the need to dissect how host genetics and microbial communities jointly shape fitness across diverse environments (McMullen et al., 2020; Uy et al., 2015). This dual capacity for genetic and microbial adaptation is especially critical as climate change and habitat degradation intensify, threatening species’ ability to cope with novel stressors (Gilbert et al., 2012; van Opstal & Bordenstein, 2015). Furthermore, it is also possible that the genetic differences of the *D. basisetae* populations alter the microbiome that further alters the adaptation of the host species. In this context, microbiome-based conservation strategies, such as augmenting beneficial microbial associations or performing targeted transplants with slurries sourced from local or related species, offer promising routes to bolster resilience (Amend et al., 2022; Mueller et al., 2020). Such approaches could be particularly impactful for Hawaiian *Drosophila*, many of which occupy narrow ecological niches and face elevated risks from climate change, invasive predators, and habitat degradation and fragmentation (Henry et al., 2013; Houwenhuyse et al., 2021; Price & Muir, 2008). For instance, rearing captive colonies with site-specific microbiomes could enhance thermal tolerance and reproductive success prior to release, improving restoration outcomes. Given the low genetic differentiation among *D. basisetae* populations, microbiome manipulation could promote local adaptation without disrupting population structure (Douglas, 2015). However, scaling up these interventions requires a better understanding of the long-term stability and heritability of introduced microbiomes, their effects on native microbial networks, and the broader ecological consequences (Erickson et al., 2004; Lynch & Hsiao, 2019; van Opstal & Bordenstein, 2015). Integrating microbiome management with habitat protection, invasive-species control, and genetic monitoring will be essential to safeguard Hawaii’s picture-winged *Drosophila* amid rapid environmental change (Koch et al., 2020; Muir & Price, 2008).

## Acknowledgements

High throughput amplicon library preparation was performed by the University of Hawaiʻi Microbial Genomics and Analytical Laboratory (MGAL: https://icemhh.pbrc.hawaii.edu/mgal/). The Vermont Advanced Computing Center (VACC; URL: https://www.uvm.edu/vacc) provided computational resources that contributed to this publication. The next-generation sequencing was performed in part, at the Vermont Integrative Genomics Resource Massively Parallel Sequencing Facility and was supported by the University of Vermont Cancer Center, Lake Champlain Cancer Research Organization, UVM College of Agriculture and Life Sciences, and the UVM Larner College of Medicine. At UNLV Franklin Ung and Destiny Johnston assisted with the phenotypic measurements and bacterial and fungal culturing; Leon Kyle Boyles assisted with the EthoVision software analysis.

## Data and resource availability

Phenotypic data will be uploaded to Dryad (https://datadryad.org/stash) and microbiome data will be uploaded to the National Microbiome Data Collaborative (https://data.microbiomedata.org/). Genomic data was submitted to the National Center for Biotechnology Information’s (NCBI) Short Read Archive (SRA). All samples are available under BioProject: **PRJNA1176294**. SRR numbers for each sample can be found in **Table S1**. The code to reproduce the genomic analyses can be found at: https://github.com/Jcbnunez/thermofly

---

**Table S1:** Metadata for genomic samples.

| accession | bioproject_accession | biosample_accession | sample_name | Site |
| --- | --- | --- | --- | --- |
| SRR31077433 | PRJNA1176294 | SAMN44386966 | D_bas.wild.US-HI-Tom.27_1_2023.w.DP_B_T_1.1 | Tom's Trail |
| SRR31077432 | PRJNA1176294 | SAMN44386967 | D_bas.wild.US-HI-Tom.27_1_2023.w.DP_B_T_2.1 | Tom's Trail |
| SRR31077421 | PRJNA1176294 | SAMN44386968 | D_bas.wild.US-HI-Tom.27_1_2023.w.DP_B_T_3.1 | Tom's Trail |
| SRR31077418 | PRJNA1176294 | SAMN44386969 | D_bas.wild.US-HI-Tom.27_1_2023.w.DP_B_T_4.1 | Tom's Trail |
| SRR31077417 | PRJNA1176294 | SAMN44386970 | D_bas.wild.US-HI-Tom.27_1_2023.w.DP_B_T_5.1 | Tom's Trail |
| SRR31077416 | PRJNA1176294 | SAMN44386971 | D_bas.wild.US-HI-Tom.27_1_2023.w.DP_B_T_6.1 | Tom's Trail |
| SRR31077415 | PRJNA1176294 | SAMN44386972 | D_bas.wild.US-HI-Tom.27_1_2023.w.DP_B_T_7.1 | Tom's Trail |
| SRR31077414 | PRJNA1176294 | SAMN44386973 | D_bas.wild.US-HI-Ola.27_1_2023.w.DP_B_O_1.1 | Ola'a |
| SRR31077413 | PRJNA1176294 | SAMN44386974 | D_bas.wild.US-HI-Ola.27_1_2023.w.DP_B_O_2.1 | Ola'a |
| SRR31077412 | PRJNA1176294 | SAMN44386975 | D_bas.wild.US-HI-Ola.27_1_2023.w.DP_B_O_3.1 | Ola'a |
| SRR31077431 | PRJNA1176294 | SAMN44386976 | D_bas.wild.US-HI-Ola.27_1_2023.w.DP_B_O_4.1 | Ola'a |
| SRR31077430 | PRJNA1176294 | SAMN44386977 | D_bas.wild.US-HI-Ola.27_1_2023.w.DP_B_O_5.1 | Ola'a |
| SRR31077429 | PRJNA1176294 | SAMN44386978 | D_bas.wild.US-HI-Ola.27_1_2023.w.DP_B_O_6.1 | Ola'a |
| SRR31077428 | PRJNA1176294 | SAMN44386979 | D_bas.wild.US-HI-Ola.27_1_2023.w.DP_B_O_7.1 | Ola'a |
| SRR31077427 | PRJNA1176294 | SAMN44386980 | D_bas.wild.US-HI-Ola.27_1_2023.w.DP_B_O_8.1 | Ola'a |
| SRR31077426 | PRJNA1176294 | SAMN44386981 | D_bas.wild.US-HI-Ola.27_1_2023.w.DP_B_O_9.1 | Ola'a |
| SRR31077425 | PRJNA1176294 | SAMN44386982 | D_bas.wild.US-HI-Ola.27_1_2023.w.DP_B_O_10.1 | Ola'a |
| SRR31077424 | PRJNA1176294 | SAMN44386983 | D_bas.wild.US-HI-Ola.27_1_2023.w.DP_B_O_11.1 | Ola'a |
| SRR31077423 | PRJNA1176294 | SAMN44386984 | D_bas.wild.US-HI-Ola.27_1_2023.w.DP_B_O_12.1 | Ola'a |
| SRR31077422 | PRJNA1176294 | SAMN44386985 | D_bas.wild.US-HI-Ola.27_1_2023.w.DP_B_O_13.1 | Ola'a |
| SRR31077420 | PRJNA1176294 | SAMN44386986 | D_bas.wild.US-HI-Ola.27_1_2023.w.DP_B_O_14.1 | Ola'a |
| SRR31077419 | PRJNA1176294 | SAMN44386987 | D_bas.wild.US-HI-Ola.27_1_2023.w.DP_B_O_15.1 | Ola'a |

**Table S2.**
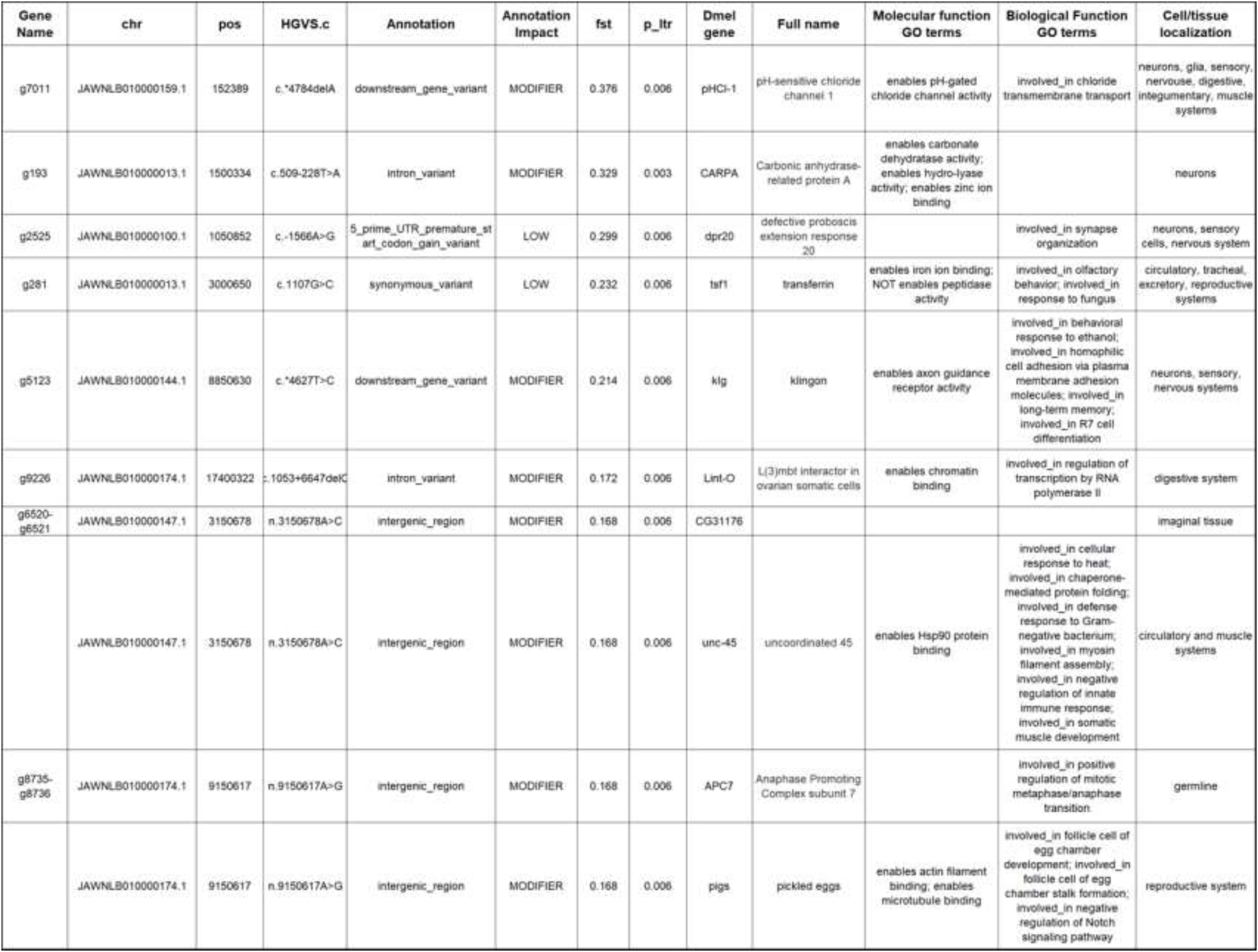
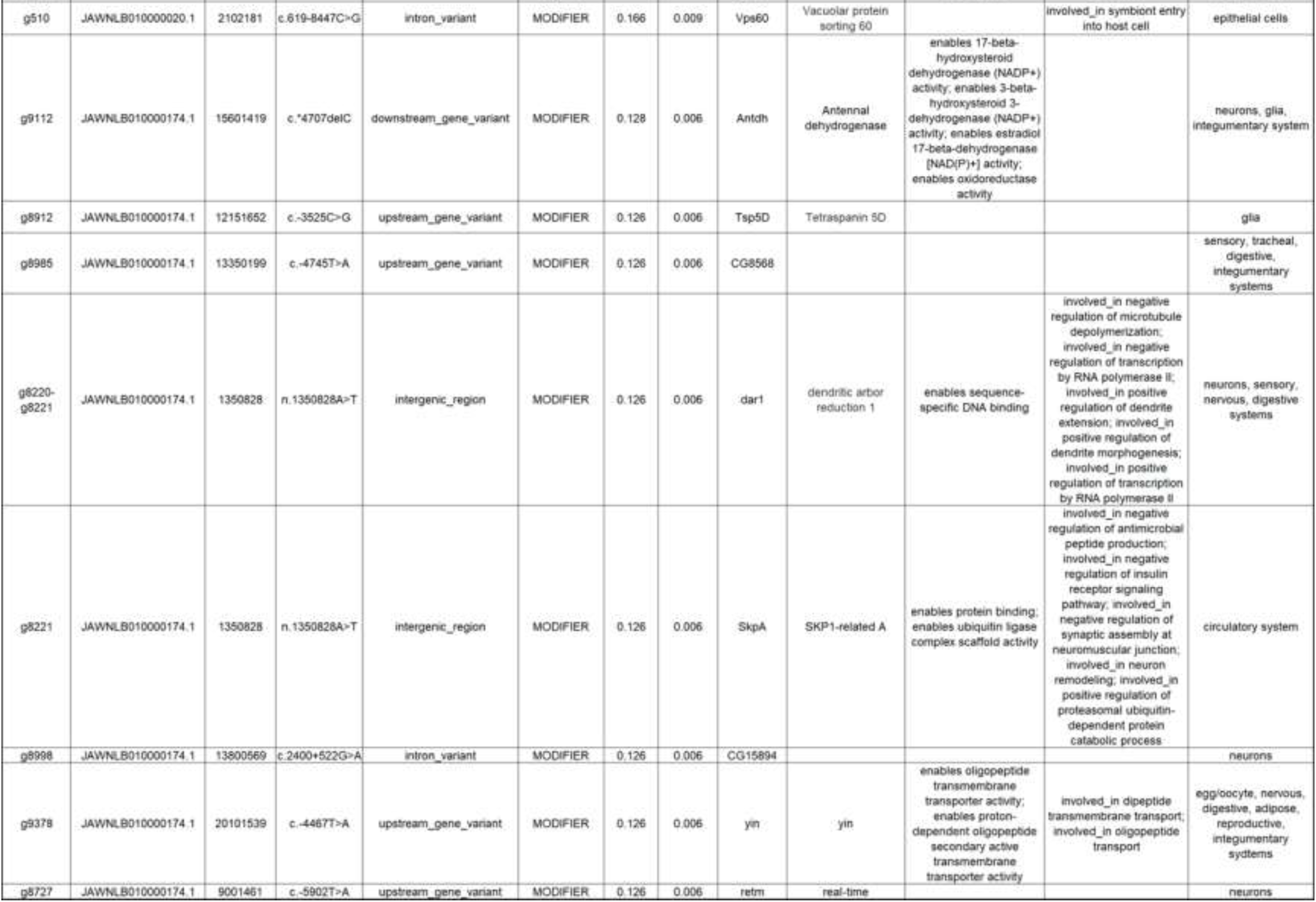
Annotation of Outlier SNPs. **Gene Name:** gene name in the ab initio prediction; **chr:** name of the scaffold; 1 **pos:** position of the SNP; **HGVS.c:** Standard HGVS Variant Nomenclature for the variant. **Annotation:** SNP annotation type inside the predicted sequence; **Impact:** predicted functional impact; **fst:** *F*ST value; **p_lrt:** *P*-value in the regression test from the likelihood ratio test. **Dmel gene:** name of the gene in *D. melanogaster*; **Full name:** full name in *D. melanogaster*; **Molecular/Biological function:** GO terms associated with gene; Cell tissue localization: based on expression from the FlyAtlas (*D. melanogaster*).

**Table S3.** Bacteria and fungi identified in the baits used to attract the flies and, in the flies, collected from both Tom’s Trail and Ola’a locations. Each unique Otu (operational taxonomic unit) is listed in rank ordered of percent abundance along with the genus name. The bacteria and fungi that were identified in the baits but not the flies are bolded and indicated by ** and the bacteria and fungi that were identified in the flies but not the baits are bolded indicated by *.

| Bacteria in Baits (n = 4) | Percent | Bacteria in flies (n = 23) | Percent |
| --- | --- | --- | --- |
| Otu002 *Leuconostoc* | 31.24 | Otu002 *Leuconostoc* | 32.16 |
| Otu016 *Pseudomonas* | 22.26 | Otu001 *Gluconobacter* | 13.47 |
| **Otu030 *Weissella* \*\*** | **12.34** | Otu007 *Gluconobacter* | 12.97 |
| Otu001 *Gluconobacter* | 7.63 | Otu016 *Pseudomonas* | 11.05 |
| **Otu037 *Lactococcus* \*\*** | **6.76** | **Otu025 *Halomonas* \*** | **6.13** |
| Otu029 *Delftia* | 5.79 | Otu011 *Acetobacter* | 4.76 |
| **Otu034 *Pseudomonas* \*\*** | **3.75** | Otu034 *Pseudomonas* | 3.87 |
| Otu007 *Gluconobacter* | 2.58 | Otu018 *Chryseobacterium* | 3.71 |
| **Otu060 *Pseudomonas* \*\*** | **2.50** | Otu029 *Delftia* | 3.63 |
| **Otu055 *Weissella* \*\*** | **2.32** | **Otu028 *Comamonas* \*** | **2.10** |
| **Otu045 *Brevundimonas* \*\*** | **1.25** | Otu022 *Stenotrophomonas* | 1.92 |
| Otu011 *Acetobacter* | 0.77 | **Otu024 *Sphingobacterium* \*** | **1.91** |
| Otu018 *Chryseobacterium* | 0.51 | **Otu060 *Pseudomonas* \*** | **1.18** |
| **Otu050 *Alcaligenes* \*\*** | **0.26** | Otu035 *Chryseobacterium* | 1.01 |
|  |  | **Otu078 *Stenotrophomonas* \*** | **0.07** |

| Fungi in baits (n=4) | Percent | Fungi in flies (n=23) | Percent |
| --- | --- | --- | --- |
| OTU97_43 *Wickerhamomyces* | 25.00 | OTU97_11 *Limtongozyma* | 21.78 |
| OTU97_33 *Wickerhamomyces* | 13.14 | OTU97_8 *Kurtzmaniella* | 11.44 |
| OTU97_8 *Kurtzmaniella* | 9.22 | OTU97_33 *Wickerhamomyces* | 10.02 |
| OTU97_11 *Limtongozyma* | 8.50 | OTU97_14 *Kurtzmaniella* | 9.59 |
| OTU97_3 *Pichia* | 7.63 | **OTU97_18 *Geotrichum* \*** | **8.31** |
| OTU97_32 *Wickerhamomyces* | 5.50 | OTU97_25 *Candida* | 5.72 |
| OTU97_4 *Pichia* | 5.48 | **OTU97_12 *Pichia* \*** | **3.25** |
| OTU97_40 *Wickerhamomyces* | 4.72 | OTU97_20 *Zygoascus* | 3.09 |
| OTU97_14 *Kurtzmaniella* | 3.55 | OTU97_19 *Wickerhamomyces* | 2.57 |
| OTU97_19 *Wickerhamomyces* | 3.14 | OTU97_3 *Pichia* | 2.55 |
| OTU97_26 *Wickerhamomyces* | 2.62 | OTU97_4 *Pichia* | 2.30 |
| OTU97_50 *Geotrichum* | 2.59 | OTU97_113 *Meyerozyma* | 2.27 |
| OTU97_6 *Pichia* | 1.57 | OTU97_43 *Wickerhamomyces* | 2.13 |
| OTU97_62 *Pichia* | 1.30 | OTU97_32 *Wickerhamomyces* | 2.09 |
| OTU97_20 *Zygoascus* | 1.05 | OTU97_6 *Pichia* | 1.73 |
| OTU97_121 *Pichia* | 1.04 | OTU97_26 *Wickerhamomyces* | 1.71 |
| OTU97_13 *Kurtzmaniella* | 0.64 | **OTU97_38 *Trigonopsis* \*** | **1.63** |
| OTU97_10 *Geotrichum* | 0.53 | OTU97_10 *Geotrichum* | 1.42 |
| OTU97_37 *Geotrichum* | 0.53 | OTU97_13 *Kurtzmaniella* | 1.31 |
| OTU97_25 *Candida* | 0.52 | OTU97_40 *Wickerhamomyces* | 1.25 |
| OTU97_31 *Candida* | 0.52 | OTU97_50 *Geotrichum* | 0.97 |
| OTU97_113 *Meyerozyma* | 0.32 | **OTU97_54 *Papiliotrema* \*** | **0.88** |
| **OTU97_15 *Penicillium* \*\*** | **0.32** | OTU97_37 *Geotrichum* | 0.87 |
| **OTU97_135 *Wickerhamomyces*\*\*** | **0.26** | OTU97_62 *Pichia* | 0.24 |
| **OTU97_67 *Saccharomycopsis* \*\*** | **0.26** | **OTU97_30 *Geotrichum* \*** | **0.19** |
|  |  | **OTU97_9 *Geotrichum* \*** | **0.18** |
|  |  | **OTU97_58 *Pichia* \*** | **0.14** |
|  |  | **OTU97_159 *Pichia* \*** | **0.10** |
|  |  | OTU97_31 *Candida* | 0.09 |
|  |  | OTU97_121 *Pichia* | 0.09 |
|  |  | **OTU97_41 *Pichia* \*** | **0.05** |
|  |  | **OTU97_55 *Pichia* \*** | **0.05** |
|  |  | **OTU97_166 *Pichia* \*** | **0.05** |

**Fig. S1.**
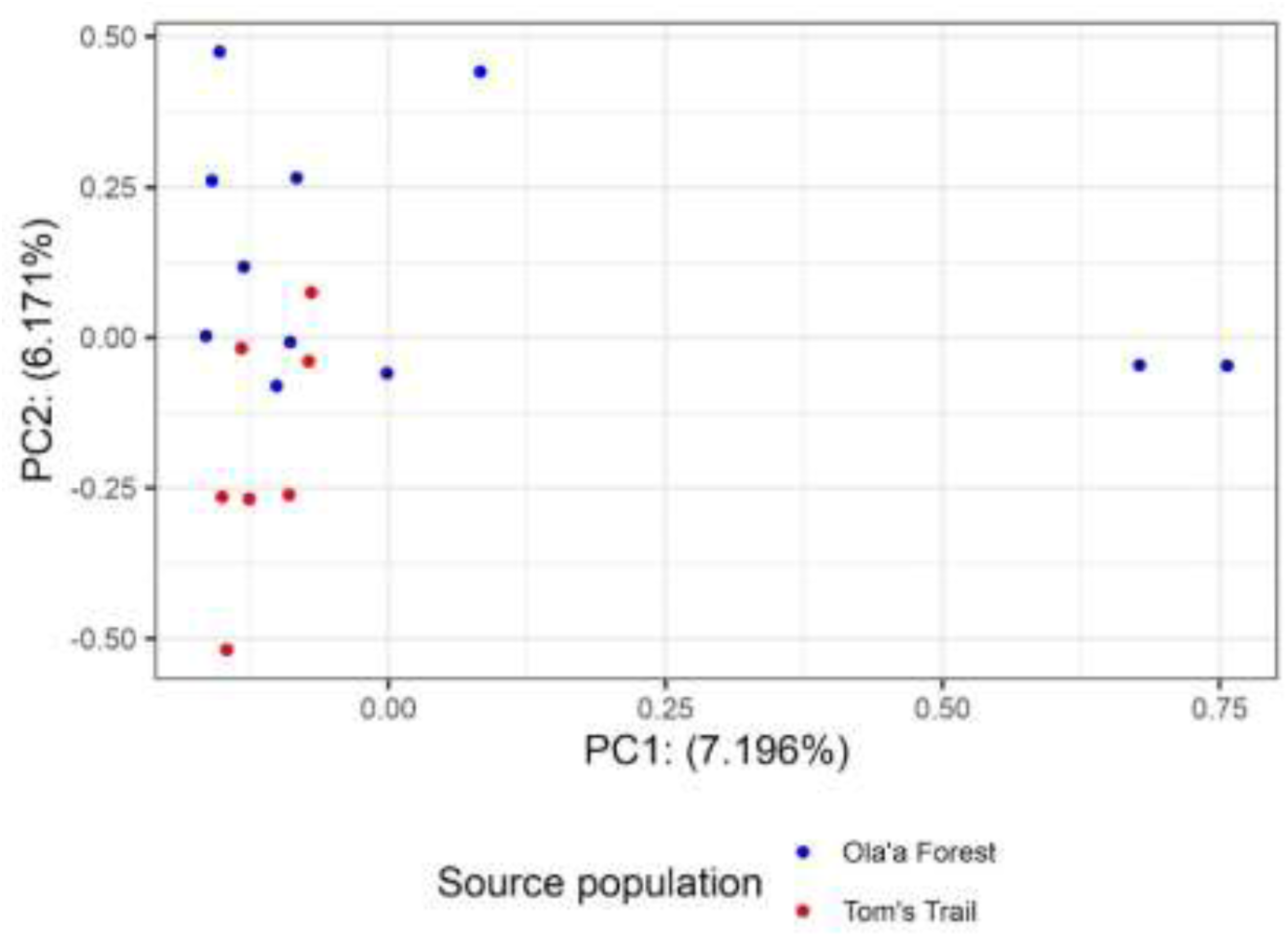
Principal component analysis on SNPs derived from the genotype likelihood pipeline.

**Fig. S2.**
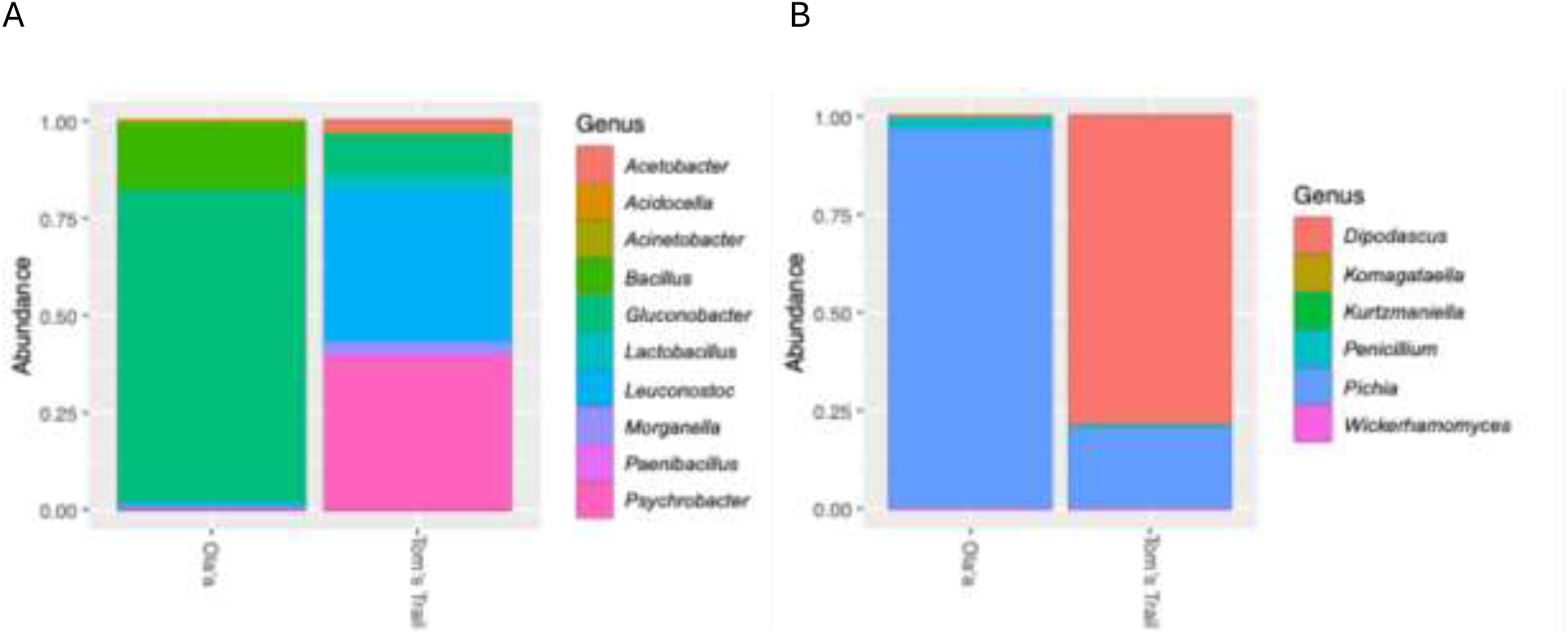
Microbiome of slurries collected from cultured feces of live *D. basisetae* flies returned to the laboratory: The relative frequencies of (A) Bacteria and (B) Fungi from Tom’s Trail (n=1 for bacteria and n=2 for fungi) & Olaʻa (n=2 for bacteria and n=3 for fungi). The relative frequencies of bacteria from Ola’a was 81% *Gluconobacter*, 17.3% *Bacillus*; the relative frequencies of bacteria from Tom’s Trails was 11% *Gluconobacter*, 40% *Leuconostoc*, 40% *Psychrobacter* 3% *Morganella* 3%, 3% *Acetobacter* 3%, and 3% *Lactobacillus*. The relative frequencies of Fungi from Ola’a was 97% *Pichia*, 3 % *Penicillium*. The relative frequencies of Fungi from Tom’s Trail was 21% *Pichia*, 79% *Dipodascus*.

**Fig. S3.**
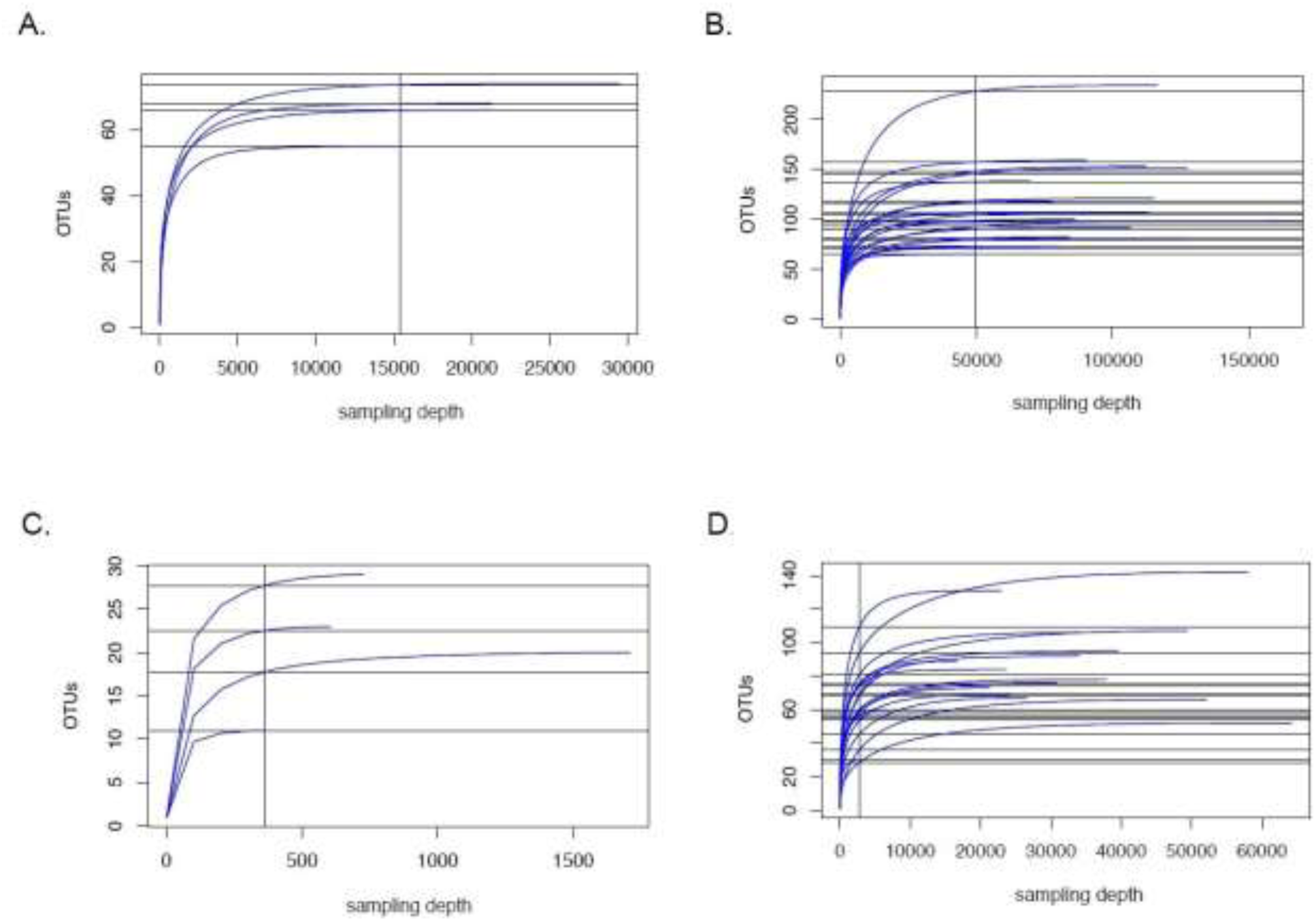
Rarefaction curves for non-subsampled read data for *D. basisetae*. A. 16S rRNA, Olaʻa, n=4. B. 16S rRNA, Tom’s Trail, n=19. C. ITS, Olaʻa, n=4. D. ITS, Tom’s Trail, n=19.

**Fig. S4.**
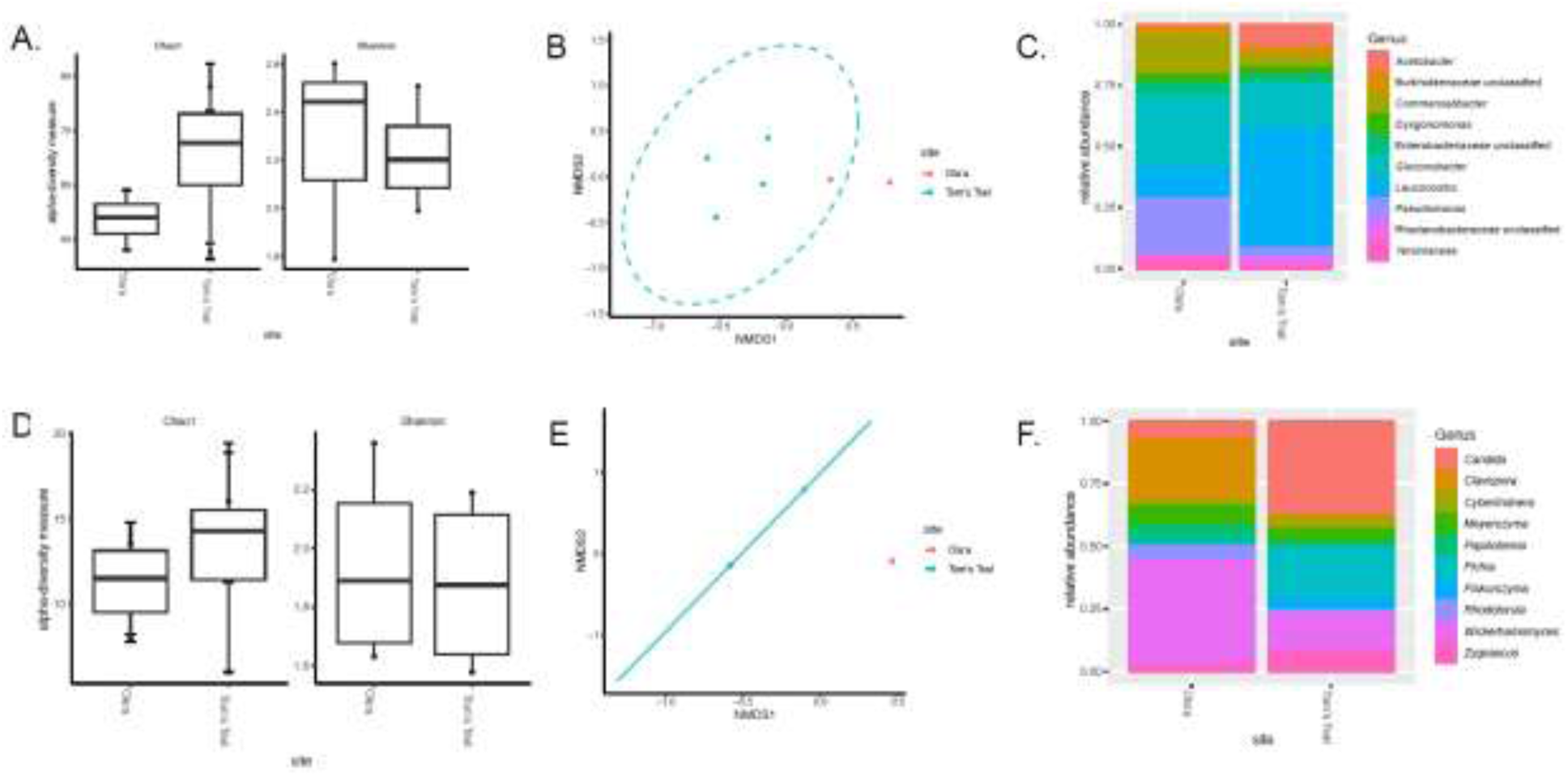
Microbiome of wild *D. basisetae* flies: Alpha and Beta Diversity measures of bacterial and fungal taxa from Tom’s Trail and Ola’a. (A) Taxonomic richness (based on Chao1) of bacterial taxa did not differ between Ola’a and Tom’s Trail (*P* = 0.4); nor did Shannon diversity did not differ between sites (*P* = 0.63). (B) Beta Diversity of bacterial taxa (based on ANOSIM using Jaccard’s distance) differed significantly between sites (*P* = 0.02). (C) The relative frequencies of the ten most common bacterial taxa in both locations; no taxa differed significantly in terms of abundance. (D) Taxonomic richness (based on Chao1) of fungal taxa was not significantly different between sites (*P* = 0.49) nor was Shannon diversity (P = 0.89). (E) Beta Diversity of fungi (based on ANOSIM using Jaccard’s distance) was significantly different between sites (*P* = 0.027). (F) The relative frequencies of the ten most common fungal taxa in both locations: *Clavispora* (*P* = 0.057), *Candida* (*P* = 0.057), and *Wickerhamomyces* (*P* = 0.057) nearly differed significantly. N=4 randomly selected flies for Tom’s Trail and n=4 for Olaʻa.

